# The ELF3-BBX24/BBX25-PIF4 module controls thermosensory growth in Arabidopsis

**DOI:** 10.1101/2024.02.14.580362

**Authors:** Bidhan Chandra Malakar, Shivani Singh, Vikas Garhwal, Rajanesh Chandramohan, Gouranga Upadhyaya, Sreeramaiah N. Gangappa

**Affiliations:** Department of Biological Sciences, Indian Institute of Science Education and Research Kolkata, Mohanpur 741246, West Bengal, India; Centre for Climate and Environmental Studies, Indian Institute of Science Education and Research Kolkata, Mohanpur 741246, West Bengal, India

**Keywords:** Thermomorphogenesis, BBX24, BBX25, PIF4, ELF3, gene regulation

## Abstract

Temperature serves as a crucial environmental cue governing the growth and adaptation of plants in their natural habitat. PHYTOCHROME INTERACTING FACTOR 4 (PIF4) is a central regulator that promotes thermomorphogenesis in Arabidopsis. Understanding its precise regulation is critical for optimal thermomorphogenic growth. Here, we identified two BBX proteins, BBX24 and BBX25, as novel components of the PIF4-mediated thermosensory pathway and act to promote warm temperature-mediated growth. The *bbx24* and *bbx25* single and double mutants showed moderate to strong temperature-insensitive hypocotyl and cotyledon growth. Warm temperature induces *BBX24* and *BBX25* mRNA expression and protein accumulation. Genetic and biochemical analysis revealed that BBX24/BBX25 promotes PIF4-mediated thermosensory growth by counteracting a key component of the evening complex, ELF3. While ELF3 inhibits *BBX24*/*BBX25* gene expression at low ambient temperatures in the evening, warm temperature-mediated inhibition of ELF3 activity results in enhanced BBX24/BBX25 activity. Moreover, BBX24/25 inhibit ELF3 function through direct physical interaction and likely relieves repression on PIF4, enhancing its activity and thermomorphogenesis. Together, this study unravels ELF3-BBX24/BBX25-PIF4 as a key regulatory module that controls growth and development under varying temperature cues.

## INTRODUCTION

Ecological cues such as light and temperature are key in regulating plant growth, architecture, and fitness (Casal and Balasubramanian, 2019; Casal and Questa, 2018; Franklin et al., 2014; Legris et al., 2019; Lippmann et al., 2019; Quint et al., 2023). Plants have evolved sophisticated and multifaceted mechanisms to perceive, integrate, and process these cues (Krahmer and Fankhauser, 2023; Legris *et al*., 2019). Because plants experience these conflicting signals, they must constantly sense and integrate them to optimise growth and acclimate to local environments. Warm ambient temperatures strongly influence plant growth by accelerating vegetative and reproductive transitions, resulting in shortened life-cycle duration (Casal and Balasubramanian, 2019; Crawford et al., 2012; Koini et al., 2009; Krahmer and Fankhauser, 2023; Kumar et al., 2012; Quint *et al*., 2023). However, warm temperatures strongly compromise immunity and severely affect seed set and yield (Burroughs et al., 2023; Koester et al., 2014; Zhao et al., 2017). Increasing seasonal growth temperature due to global warming impairs plant growth and yield, threatening farmers’ livelihood and food security (Peng et al., 2004; Zhao *et al*., 2017). Therefore, understanding how plants sense and integrate temperature cues is critical to delineating thermosensory signalling mechanisms to breed climate-resilient crops.

The light signaling pathway consists of photoreceptors and downstream signaling molecules that have been shown to play a pivotal role in sensing and integrating temperature cues (Legris *et al*., 2019; Li et al., 2016). Key studies revealed that red-light absorbing phytochrome B (phyB) senses temperature cues (Jung et al., 2016; Legris et al., 2016). PhyB exists in two interconvertible physiologically active (Pfr) and inactive (Pr) forms. Warm temperature favours the conversion of Pfr to Pr form, reducing the phyB activity (Legris *et al*., 2016). The phyB targets a group of growth-promoting transcription factors, PHYTOCHROME INTERACTING FACTORS (PIFs), for phosphorylation and degradation in a red-light-dependent manner (de Lucas et al., 2008; Huq and Quail, 2002; Lorrain et al., 2008). Among PIFs, PIF4 and PIF7 have been shown to mediate thermomorphogenesis, respectively, under SD and LD photoperiods (Burko et al., 2022; Chung et al., 2020; Delker et al., 2022; Fiorucci et al., 2020; Koini *et al*., 2009; Kumar *et al*., 2012). PIF4 is essential for warm temperature-induced hypocotyl and petiole growth, rosette development, hyponasty, and flowering (Casal and Balasubramanian, 2019; Favero et al., 2020; Koini *et al*., 2009; Kumar *et al*., 2012; Oh et al., 2012; Quint *et al*., 2023; Quint et al., 2016; Zhao and Bao, 2021). PIF4-mediated thermomorphogenesis is linked to the activation of genes involved in auxin/brassinosteroid biosynthesis and signaling (Franklin et al., 2011; Ibanez et al., 2018; Krahmer and Fankhauser, 2023; Sun et al., 2012), cell elongation (Gangappa and Kumar, 2017) and flowering(Kumar *et al*., 2012). Warm temperature-mediated inhibition of phyB activity increases PIF4 function, resulting in thermosensory growth (Legris *et al*., 2016). Unlike phyB, the blue light receptor Cryptochrome 1 (CRY1) inhibits thermomorphogenesis through PIF4 sequestration(Ma et al., 2016). The central oscillator components of the circadian clock, including the evening complex (EC), play a crucial role in controlling thermomorphogenesis by regulating PIF4 function. The EARLY FLOWERING 3 (ELF3), a critical component of the evening complex (EC) and a thermosensor, inhibits *PIF4* transcription and also sequesters it by forming heterodimers (Box et al., 2015; Jung et al., 2020; Nieto et al., 2015; Nusinow et al., 2011). Moreover, upstream regulatory components such as CONSTITUTIVE PHOTOMORPHOGENIC 1 (COP1), DE-ETIOLATED 1 (DET1) and SUPPRESSOR OF PHYTOCHROME A-105 1 (SPA1) proteins promote PIF4-mediated thermomorphogenic growth by stabilizing PIF4 protein (Gangappa and Kumar, 2017; Lee et al., 2020; Park et al., 2017). Recent studies also reveal that the HEMERA/REGULATOR OF the CHLOROPLAST BIOGENESIS (HMR/RCB) module works cooperatively to stabilize PIF4 during the day and promote thermomorphogenesis (Qiu et al., 2019; Qiu et al., 2021). The Heat shock transcription factor (HSF) family members, HSFA1a/b/d/e, interact with PIF4 and stabilize it during light by interfering with PIF4-phyB interaction, resulting in thermomorphogenesis during the day (Tan et al., 2023a).

The B-BOX containing zinc-finger (BBX) proteins are highly conserved transcription factors across eukaryotes that regulate gene expression by directly binding to DNA or as transcriptional co-activators or co-repressors through protein-protein interaction. BBX proteins have emerged as crucial integrators of various environmental cues (Ding et al., 2018; Gangappa and Botto, 2014; Khanna et al., 2009; Song et al., 2020; Vaishak et al., 2019; Yuan et al., 2021). The BBX24 and BBX25 are close homologs that function as transcriptional co-repressors of HY5 to inhibit photomorphogenesis (Gangappa et al., 2013; Job et al., 2018). Additionally, BBX24 functions in various signaling pathways, including UV-B signaling (Huang et al., 2022; Jiang et al., 2012), shade-avoidance response (Crocco et al., 2015; Saura-Sanchez et al., 2023) and flowering (Li et al., 2014). However, the role of BBX24 and BBX25 in temperature-mediated regulation of growth and development remains unknown. Here, we report that BBX24/BBX25 as key regulators of PIF4-mediated thermosensory growth. We found that *BBX24*/*BBX25* gene expression and protein levels are enhanced at warm temperatures, further enhancing PIF4 protein accumulation and thermomorphogenesis. Our study demonstrates that ELF3 functions upstream of BBX24/BBX25, inhibiting their gene expression during the evening hours under low temperatures. However, warm temperature-mediated inhibition of ELF3 activity results in increased BBX24/BBX25 function and, in turn, enhances PIF4-mediated thermomorphogenic growth. Additionally, BBX24/BBX25 inhibit ELF3 protein accumulation resulting in the release of the negative inhibitory effect of ELF3 on PIF4 protein and transcript accumulation. Together, this study uncovers a novel regulatory mechanism by which BBX24/BBX25 promote PIF4-mediated thermosensory growth via antagonizing ELF3 function.

## RESULTS

### BBX24 and BBX25 are essential for warm temperature-mediated growth

To unravel the possible role of BBX24 and BBX25 in temperature-mediated regulation of hypocotyl growth, we analyzed the hypocotyl growth response of wild-type (Col-0) and *bbx24* and *bbx25* mutants grown under 22°C and 27°C. Our data revealed that *bbx24* and *bbx25* had moderate but significantly shorter hypocotyls at 22°C and 27°C than Col-0 both under short-day (SD) and long-day (LD) photoperiods (Figure 1A-1C). Compared to Col-0, *bbx24* showed stronger inhibition in hypocotyl growth than the *bbx25* mutant at 27°C (Figure 1A-1C). Interestingly, the *bbx24bbx25* double mutant had a much shorter hypocotyl length at 27°C than the single mutants (Figure 1A-1C), suggesting their additive function in promoting warm temperature-mediated hypocotyl growth. Contrary to mutants, the *BBX24* overexpression (hereafter referred to as *BBX24-OE*) transgenic line showed enhanced thermosensory hypocotyl growth as they had significantly longer hypocotyls than Col-0 at 22°C and 27°C both under SD and LD (Figure 1D-1F). To further understand whether overexpression of *BBX25* has a similar effect to *BBX24-OE* in promoting hypocotyl growth, we generated multiple transgenic lines overexpressing *BBX25* (Supplemental Figure S1A and 1B). Like *BBX24-OE* lines, transgenic lines overexpressing *BBX25* showed exaggerated hypocotyl growth compared to Col-0 at 22°C and 27°C, under SD and LD (Figure 1D-1F; Supplemental Figure S1C-S1E). Warm temperature-mediated hypocotyl elongation is attributed to increased cell elongation, which is due to increased growth-promoting hormones such as auxin and brassinosteroid (Franklin *et al*., 2011; Ibanez *et al*., 2018; Oh *et al*., 2012). To know if BBX24/BBX25-mediated hypocotyl growth is due to increased cell elongation, we analyzed the cell length of hypocotyl epidermal cells from seedlings grown at 22°C and 27°C under SD. Our data reveal that the hypocotyl epidermal cell length was significantly longer in Col-0 at 27°C than at 22°C (Figure 1G). However, in the *bbx24bbx25* double mutant, the cell length was strongly inhibited (Figure 1G). Contrarily, *BBX24* and *BBX25* overexpression lines had significantly longer cells at both 22°C and 27°C than Col-0 (Figure 1G). Together, these results suggest that BBX24/BBX25-mediated hypocotyl growth in response to warm temperature is linked to an increase in cell elongation of hypocotyl epidermal cells.

**Figure 1.**
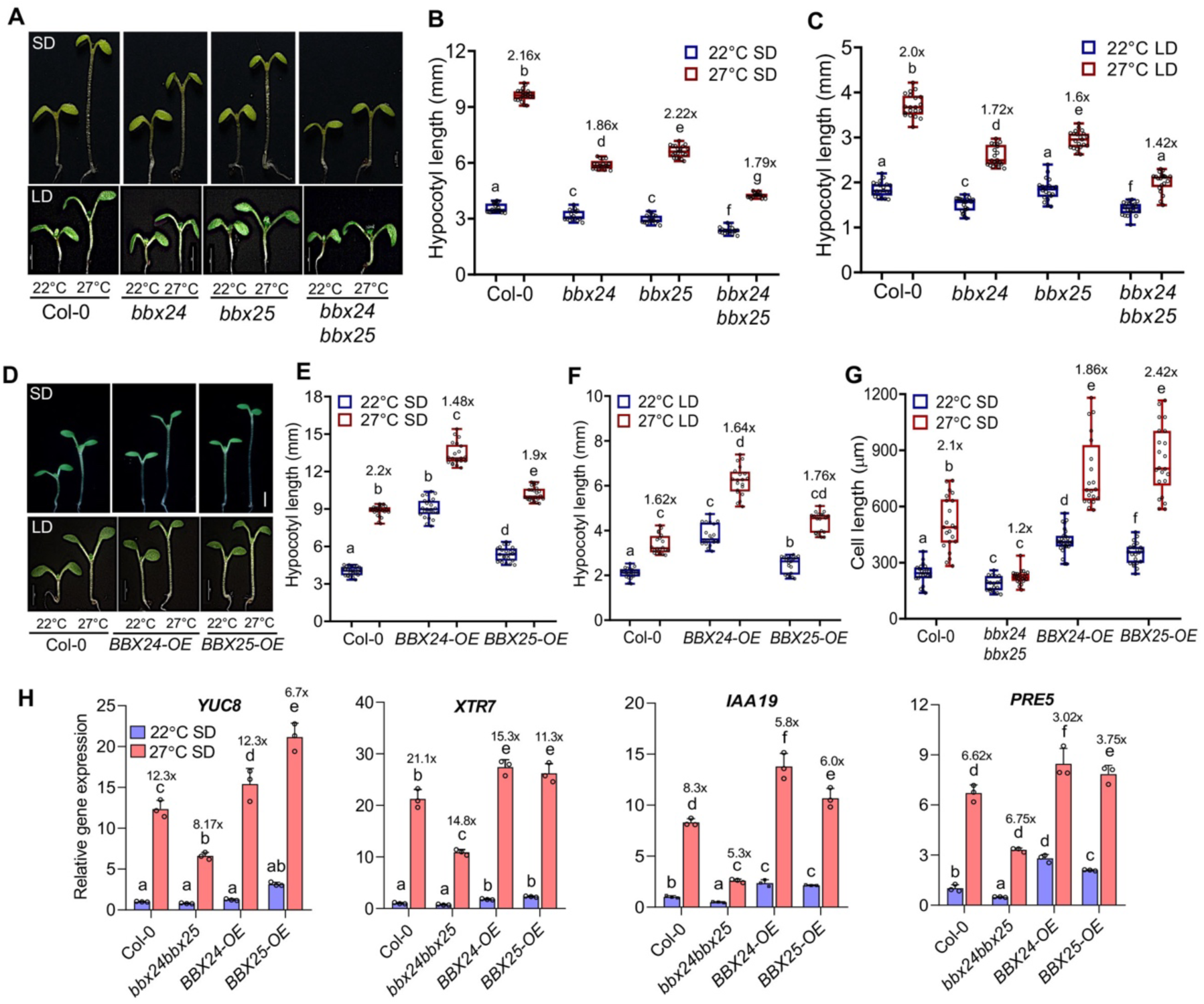
BBX24 and BBX25 promote thermosensory hypocotyl growth and gene expression. **(A)** Representative images of six-day-old Col-0, *bbx24*, *bbx25*, and *bbx24bbx25* seedlings grown at 22°C and 27°C under short-day (SD, upper panel) and long-day (LD, lower panel) photoperiods. **(B-C)** Hypocotyl length measurements (mean ± SD; n ≥ 20) of SD (**b**) and LD (**c**) grown seedlings that are shown in **A**. **(D)** Representative seedling images of Col-0, *BBX24-OE*, and *BBX25-OE* lines (line #16) were grown at 22°C and 27°C for six days under SD (upper panel) and LD (lower panel) conditions. **(E** and **F)** Hypocotyl length (mean ± SD; n ≥ 20) of SD (**E**) and LD (**F**) grown seedlings shown in (**D**). **(G)** Cell length (mean ± SD; n ≥ 15) of hypocotyl epidermal cells from indicated genotypes grown at 22°C and 27°C for six days under SD. **(H)** qRT-PCR analysis (mean ± SD of three biological replicates) showing relative transcript levels of thermoresponsive genes *YUC8*, *XTR7*, *IAA19* and *PRE5* in the indicated genotypes that were grown for six days at 22°C and 27°C under SD, and the samples were harvested at ZT23. *EF1α* was used as an internal control. In Figs. B, C, E-**G**, and **h**, different lowercase letters, shown above the bars, denote statistically significant differences between the samples (Two-way ANOVA followed by Tukey’s post-hoc test, p < 0.05). Numbers indicate fold changes between 22°C and 27°C. All the experiments were repeated at least three times, and similar results were obtained. In Figure **A** and **D**, the scale bar represents 1 mm and 2 mm, respectively. All the experiments were repeated three times, and similar results were observed.

Warm temperature-mediated hypocotyl growth is linked to enhanced accumulation of auxin and brassinosteroid (BR) growth hormones (Franklin *et al*., 2011; Gray et al., 1998; Ibanez *et al*., 2018; Krahmer and Fankhauser, 2023; Oh *et al*., 2012; Stavang et al., 2009). As *bbx24bbx25* double mutant has extremely short hypocotyls under warm temperatures, we wanted to see if exogenous application of auxin or BR can rescue this phenotype. Our data suggest that in response to a synthetic auxin picloram (PIC) or epibrassinolide (eBL) the short hypocotyl phenotype of *bbx24bbx25* was partially recovered with a higher % hypocotyl growth at 27°C compared to Col-0 (Supplemental Figure S2A-F). In contrast, *BBX24-OE* and *BBX25-OE* lines showed enhanced sensitivity to both the hormones with a reduced % hypocotyl growth compared to Col-0 (Supplemental Figure S2A-F). Moreover, we also found that overexpression of *BBX24* and *BBX25* enhanced the promoter activity of *DR5*, a synthetic auxin reporter (Supplemental Figure S2G). Contrastingly, when we treated the *BBX24-OE* and *BBX25-OE* seedlings with auxin inhibitor 2,3,5-triiodobenzoic acid (TIBA, an auxin transport inhibitor) and a BR biosynthetic inhibitor, Propiconazole (PPZ), they showed enhanced sensitivity to the inhibitor similar to Col-0 (Supplemental Figure S3A-F). Thus, these results suggest that BBX24/BBX25 indeed promote auxin/BR governed hypocotyl growth in response to warmth.

### BBX24/BBX25 function is necessary for the thermoresponsive gene expression

Consistent with the hypocotyl data, our qRT-PCR results from SD-grown seedlings harvested at ZT23 revealed that auxin biosynthetic gene *YUCCA8* (*YUC8*), cell-wall elongation gene *XYLOGLUCAN ENDOTRANSGLYCOSYLASE 7* (*XTR7*), auxin signaling gene *INDOLE ACETIC ACID19* (*IAA19*), BR signaling gene *PACLOBUTRAZOL RESISTANCE5* (*PRE5*) showed significant downregulation in the *bbx24bbx25* mutant at 27°C, while moderately reduced at 22°C, compared to Col-0 (Figure 1H). However, in the *BBX24* and *BBX25* overexpression lines, these genes showed significant upregulation both at 22°C and 27°C (Figure 1H). Additionally, our qPCR data from SD-grown seedlings harvested during daytime at ZT3 revealed that expression of growth and hormone signaling genes such as *YUC8*, *DWARF4* (*DWF4*), *Gibberellin 3-oxidases* (*GA3OX1*), *IAA19*, *PRE5*, *SMALL* AUXIN *UP-REGULATED RNA 21* (*SAUR21*), and genes involved in cell elongation, such as *EXPANSIN 8* (*EXP8*) and *XTR7* showed significant downregulation in the *bbx24bbx25* double mutant while upregulated in the overexpression lines than Col-0, at 22°C and 27°C (Supplemental Figure S4A and B). Together, these data confirm that BBX24/BBX25 induce the expression of growth-related genes in response to warm temperatures both at night and during the day.

### BBX24/BBX25 gene expression and protein levels are enhanced in response to warm temperatures

As BBX24/BBX25 promote warm temperature-mediated hypocotyl growth, we wanted to see if warm temperature regulates their gene expression and protein abundance. The qRT-PCR data from six-day-old seedlings revealed that *BBX24* and *BBX25* transcript levels showed significant upregulation at 27°C compared to 22°C both at the end of the night (ZT23) and during daytime (ZT3) (Supplemental Figure S5A). Further, when we monitored the expression of *BBX24* and *BBX25* for an extended period during the day into early night, we found that *BBX24* and *BBX25* expressions peak early in the day but their expression reduced around evening and early night, both at 22°C and 27°C (Figure 2A). Interestingly, their expression is significantly upregulated at 27°C than at 22°C both under SD (Figure 2A) and LD (Supplemental Figure S5B and C) conditions. Moreover, immunoblot analysis of the *35S::BBX24-GFP* line revealed that BBX24 protein levels was elevated at 27°C compared to 22°C, specifically at ZT2 (Supplemental Figure S5D). Further, when we monitored BBX24 protein accumulation at different ZT times, we found that at ZT0, BBX24 protein was hardly detectable, and the protein stabilized in response to light and reached the peak at ZT2 and then reduced at ZT4 and ZT8 at 22°C (Figure 2B). However, at 27°C, BBX24 protein abundance increased nearly two-fold over 22°C at ZT2, and the protein level was largely comparable between 22°C and 27°C at ZT4 and ZT8 (Figure 2B). Similarly, immunoblot analysis of the *35S::Myc-BBX25* transgenic line revealed that at ZT2, BBX25 protein levels was also elevated at 27°C compared to 22°C under SD (Supplemental Figure S5E). Specifically, BBX25 protein was more stable at ZT2 and ZT4 at 27°C compared to 22°C (Figure 2C). Unlike BBX24, BBX25 was detectable at the end of the night (ZT23), but its levels was not altered at 27°C (Figure 2C). Consistent with the increased accumulation of BBX24 protein in response to 27°C, microscopic analysis of *35S:BBX24:GFP* seedlings revealed that BBX24-GFP was more prominent in the hypocotyl region as relative fluorescence of GFP was significantly increased at 27°C than at 22°C (Figure 2D-2E), while no fluorescence was detected in Col-0 (Supplemental Figure S5F). Subsequently, we also observed an increased number of cells with BBX24-GFP fluorescence in seedlings grown at 27°C compared to 22°C (Figure 2F). Together, these results suggest that warm temperatures promote BBX24 and BBX25 gene expression and protein accumulation.

**Figure 2.**
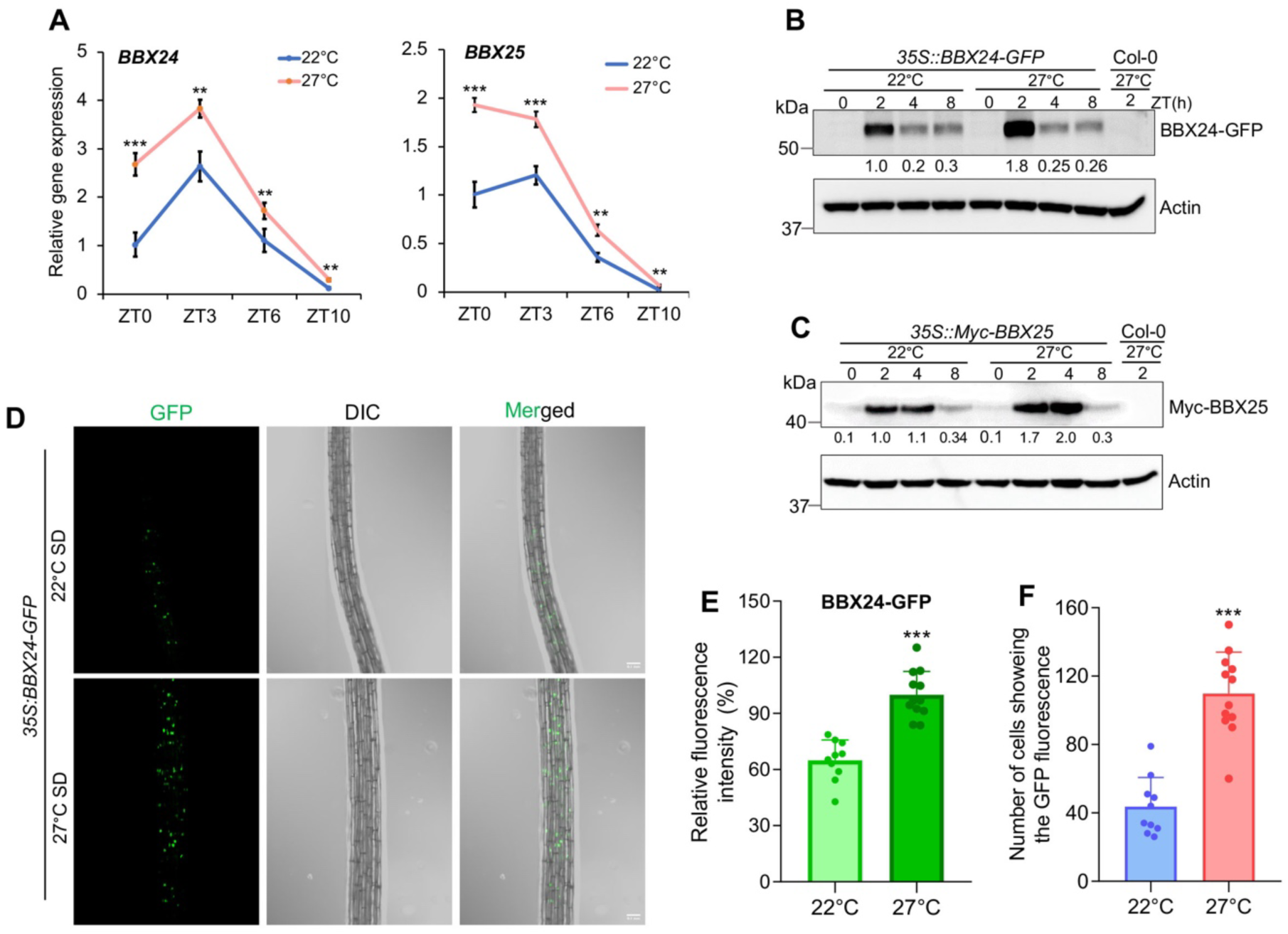
Warm temperatures promote BBX24/BBX25 transcript and protein accumulation. **(A)** qRT-PCR analysis shows *BBX24* and *BBX25* expression in six-day-old Col-0 seedlings grown at 22°C and 27°C under SD conditions, and samples were collected at indicated time points. *EF1α* was used as a loading control, and the transcript level of each sample was normalized to Col-0 22°C ZT0 time point. Error bars depict the mean ± SD of three biological replicates. Asterisks denote statistically significant differences as determined by Student’s t-test, **P<0.01, ***P<0.001. **(B)** Immunoblots showing BBX24-GFP protein level in six-days-old *35S::BBX24-GFP* seedlings grown at 22°C and 27°C under SD photoperiod and samples were harvested at different time points as indicated. **(C)** Immunoblot showing Myc-BBX25 protein status in six-days-old *35S::Myc-BBX25* seedlings grown at 22°C and 27°C under SD, and samples were harvested at different time points as indicated. In Figure **B** and **C**, 27°C grown Col-0 seedlings harvested at ZT2 were used as a negative control. Actin served as a loading control and was used for normalization. Relative protein levels (shown underneath each immunoblot) were calculated by setting 22°C grown Col-0 ZT2 to 1. The experiments were repeated three times, and similar results were obtained. **(D**-**F)** Representative images (**D**), relative GFP fluorescence intensity (**E**) and number of cells showing the fluorescence (**F**) in hypocotyls of *35S:BBX24-GFP* transgenic seedlings grown for six days after germination at 22°C and 27°C under short-day conditions. At least ten seedlings have been captured for the GFP fluorescence. Asterisks denote statistically significant differences as determined by Student’s t-test, ***P<0.001. All the experiments were repeated three times, and similar results were observed.

### BBX24/BBX25 are essential for the accumulation of PIF4 protein in response to warm temperature

As our above results indicate that BBX24/BBX25 are required for the optimal expression of growth and hormone-related genes in response to warm temperatures, we wanted to know if this is due to differential PIF4 protein abundance. Our immunoblot data from SD-grown seedlings at 22°C revealed that at the end of the night (ZT23), PIF4 protein levels was reduced in the *bbx24bbx25* double mutant than Col-0, while during the day time it was more or less similar to Col-0 (Figure 3A). Interestingly, at 27°C, the PIF4 protein levels was strongly reduced in the *bbx24bbx25* double mutant at the end of the night (ZT23) and during the day (ZT2-ZT8) compared to Col-0 (Figure 3A). However, in Col-0, PIF4 levels stayed very high at the end of the night (ZT23) and throughout the day (ZT2-ZT8) (Figure 3A). Supporting this, when 22°C grown seedlings under SD were treated with 27°C for short durations (0, 2, 4, 8h), the PIF4 protein levels in Col-0 increased with the duration of warmth treatment (Figure 3B). However, in the *bbx24bbx25*, PIF4 fails to accumulate in response to warm and its levels stayed very low than Col-0 (Figure 3B). Consistently, overexpression of *BBX24* and *BBX25* also resulted in increase in PIF4 protein levels at 22°C and 27°C compared to Col-0 (Figure 3C). BBX24/BBX25 was shown to inhibit photomorphogenesis by interfering with the HY5 transcriptional activity (Gangappa *et al*., 2013). Therefore, we wanted to see if BBX24/BBX25-mediated thermosensory hypocotyl growth is due to altered HY5 protein status. Our immunoblot data suggests that HY5 protein levels were comparable between *bbx24bbx25* double mutant and Col-0 both at 22°C and 27°C (Figure 3D), confirming that BBX24/BBX25-mediated thermosensory hypocotyl growth is not due to reduced HY5 levels but due to enhanced PIF4 protein accumulation. Together, these results suggest that the BBX24/BBX25 function is crucial for promoting PIF4 protein abundance in response to warmth both during night and daytime.

**Figure 3.**
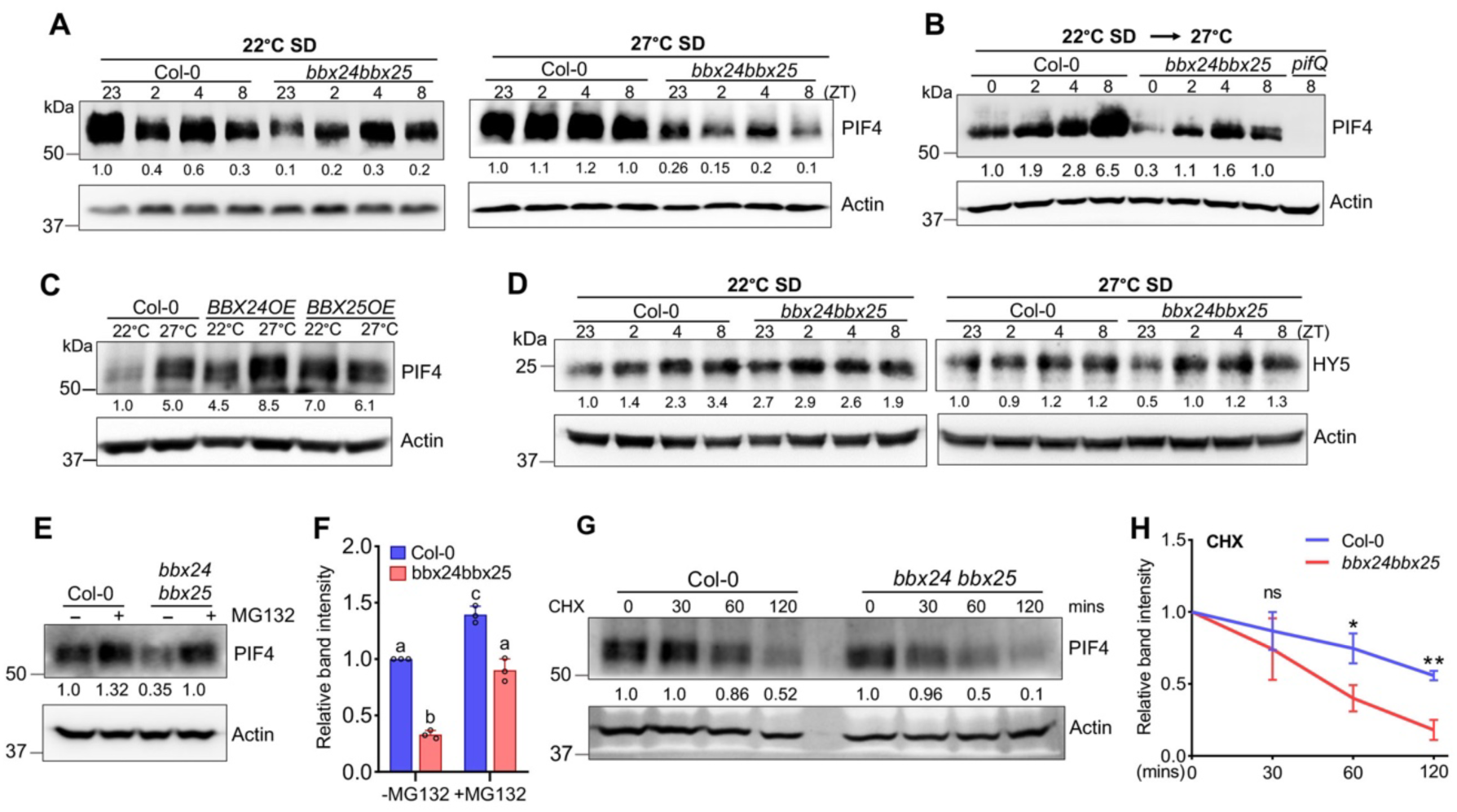
BBX24/BBX25 promotes the accumulation of PIF4 protein in response to warm temperatures. **(A)** Immunoblot analysis of PIF4 protein levels in six-day-old Col-0 and *bbx24bbx25* seedlings grown diurnally at 22°C and 27°C under SD and samples were harvested at the indicated time (ZT) points. **(B)** Immunoblot showing PIF4 protein levels in Col-0 and *bbx24bbx25* double mutant seedlings grown at 22°C SD for six days or 22°C SD grown seedlings exposed to 27°C for indicated time points. The *pifQ* mutant was used as a negative control. **(C)** Immunoblot analysis of PIF4 protein level in six-day-old Col-0, *BBX24-OE*, and *BBX25-OE* seedlings grown at constant 22°C and 27°C under SD photoperiod, and tissue were harvested at ZT23. **(D)** Immunoblot analysis of HY5 protein levels in six-day-old Col-0 and *bbx24bbx25* seedlings grown diurnally at 22°C and 27°C under SD and sampled at the indicated time (ZT) points. **(E** and **F)** Immunoblot analysis (**E**) and relative band intensity of PIF4 protein levels (**F**) in six days-old 22°C SD grown Col-0 and *bbx24bbx25* double mutant seedlings shifted to 27°C for 24hrs with MG132 treatment for 16hrs before tissue harvest. Different lowercase letters, shown above the bars, denote statistically significant differences between the samples (Two-way ANOVA followed by Tukey’s post-hoc test, p < 0.05). **(G** and **H)** Western blot analysis (**G**) and relative band intensity of PIF4 protein degradation (**H**) in six-day-old 22°C SD grown Col-0 and *bbx24bbx25* double mutants seedlings treated with 50µM MG132 before being shifted to 27°C for 8hrs were washed five times and then transferred to liquid MS medium containing 200µM CHX following tissue harvest at the indicated time points. Asterisks represent statistically significant differences determined by the Student’s t-test, *P<0.05, **P<0.01. Actin was used as a loading control in all the immunoblots. The relative PIF4 or HY5 protein level normalized to actin was shown underneath the respective immunoblot. All the experiments were repeated three times, and similar results were observed.

Next, we wanted to see if BBX24/BBX25 also affects *PIF4* at the transcriptional level; we checked its transcript levels and found that *PIF4* expression is moderately but significantly downregulated in the *bbx24bbx25* mutant while upregulated in *BBX24* and *BBX25* overexpression transgenic lines at the end of the night (ZT23), both at 22°C and 27°C under SD (Supplemental Figure S6A). However, the regulation of *PIF4* gene expression by BBX24/BBX25 is likely indirect, as our ChIP analysis followed by qPCR data revealed that BBX24/BBX25 did not bind to the *PIF4* promoter (Supplemental Figure S6B-D). Furthermore, ChIP analysis also indicates that BBX24/BBX25 do not bind to the promoters of PIF4 targeted growth-responsive genes (Supplemental Figure S6E-G).

Further, when we treated five-day-old seedlings with proteasomal inhibitor MG132, we found that protein stability increased in the *bbx24bbx25* double mutant compared to mock-treated seedlings (Figure 3E). However, it was still visibly reduced compared to MG132-treated Col-0 seedlings (Figure 3E and 3F). Consistent with this, when we treated seedlings with a translation inhibitor, cycloheximide (CHX), we found that PIF4 degradation was significantly faster in the *bbx24bbx25* mutant, specifically at 60 and 120 min of CHX treatment, compared to Col-0 (Figure 3G and 3H). Together, these data suggest that BBX24 and BBX25 play a crucial role in *PIF4* transcript accumulation and also in maintaining its protein abundance at the posttranslational level.

### BBX24/BBX25-mediated hypocotyl elongation in response to warmth is dependent on PIF4

As BBX24/BBX25 mediated warm temperature response is linked to enhanced PIF4 protein accumulation, we wanted to check the genetic interaction of BBX24/BBX25 with PIF4. We introduced *pif4* mutation into *BBX24-OE* and *BBX25-OE* lines through genetic crosses and generated homozygous lines. Analysis of hypocotyl length revealed that the exaggerated hypocotyl growth of the *BBX24-OE* line was strongly suppressed in the *pif4* mutant background as the hypocotyl length of *BBX24-OE pif4* was close to the *pif4* mutant (Supplemental Figure S7A and B). Similarly, the *BBX25-OE pif4* hypocotyl growth was markedly suppressed compared to the *BBX25-OE* line (Supplemental Figure S7A and B). These results confirm that BBX24/BBX25-mediated thermosensory hypocotyl growth strongly depends on the PIF4, implying that BBX24/BBX25 probably acts upstream of PIF4.

### BBX24/BBX25 function is essential for PIF4-mediated thermosensory growth

Consistent with the above results that BBX24/BBX25 are critical for maintaining PIF4 protein abundance, our genetic analysis also revealed that BBX24/BBX25 are necessary for PIF4-mediated thermosensory growth (Figure 4). We have introduced *bbx24*, *bbx25* and *bbx24bbx25* mutations in the *pPIF4:PIF4-FLAG* transgenic line, which overexpresses PIF4 (Gangappa et al., 2017) (hereafter referred to as *PIF4-* OE) (Supplemental Figure S8A) and analyzed their hypocotyl phenotypes. The *bbx24* or *bbx25* single mutants showed moderate suppression, while the *bbx24bbx25* strongly suppressed the exaggerated hypocotyl and cotyledon phenotype of the *PIF4-OE* line at 22°C and 27°C under SD (Figure 4A-4D; Supplemental Figure S8B-E). Similar to SD, under LD 22°C, *bbx24* and *bbx25* mutations showed additive effects in suppressing the cotyledon phenotype of the *PIF4-OE* line (Supplemental Figure S8F and G). Consistent with the growth suppression, *bbx24bbx25* strongly suppressed the expression of growth-promoting genes such as *YUC8*, *IAA19*, *XTR7*, *SAUR22* and *PRE5* in the *PIF4-OE* line (Supplemental Figure S9A).

**Figure 4.**
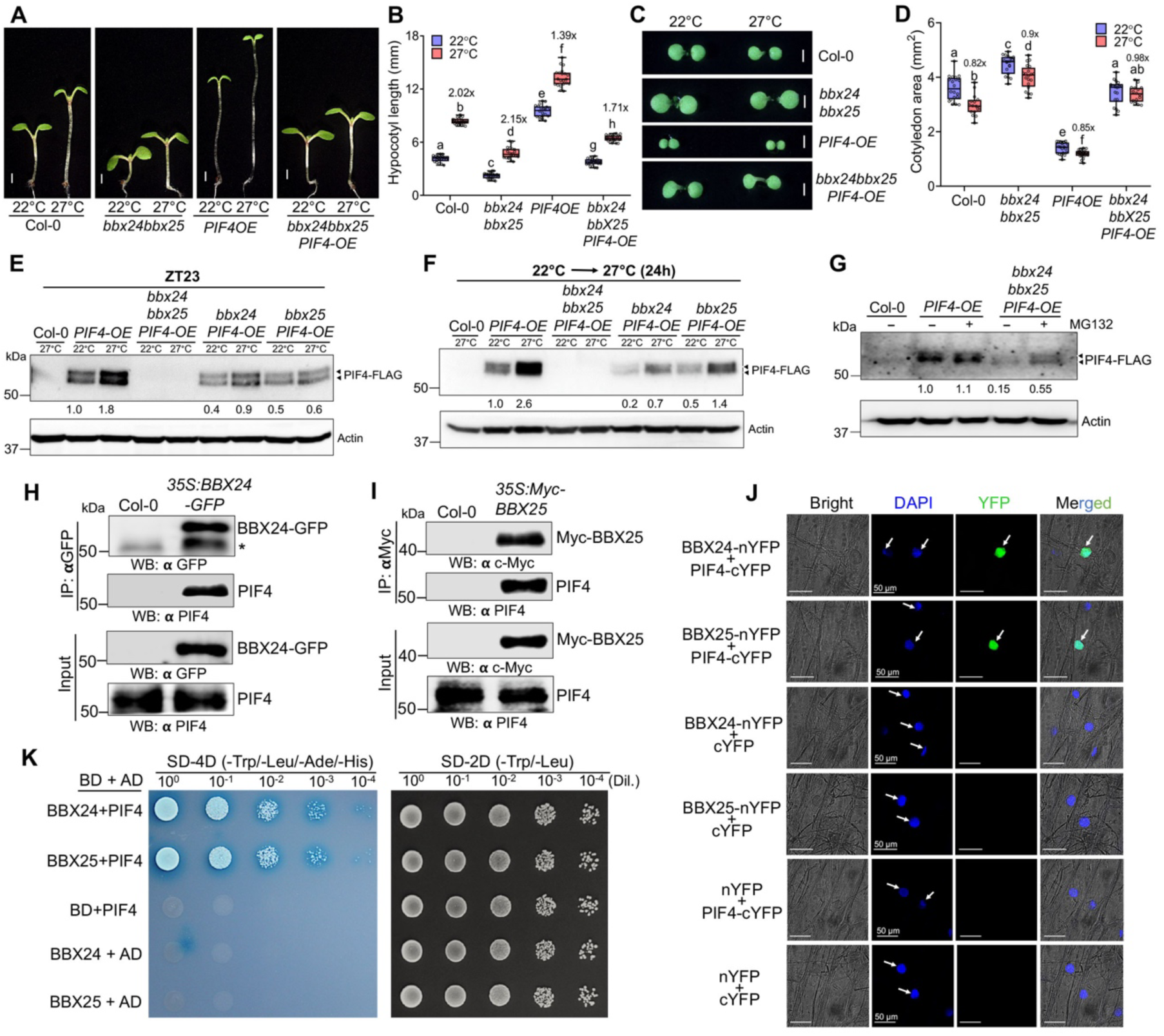
BBX24/BBX25 are essential for PIF4-mediated hypocotyl elongation in response to warm ambient temperatures. **(A** and **B)** Representative seedlings images (a) and hypocotyl length (b) of six-days-old Col-0*, bbx24bbx25, PIF4-OE*, and *bbx24bbx25PIF4-OE* seedlings grown at 22°C and 27°C under SD. The data shown is mean ± SD (n ≥ 20) seedlings. The scale bar represents 1 mm. **(C** and **D)** Representative cotyledon images (c) and cotyledon area (d) of six-day-old indicated genotypes grown at 22°C and 27°C under SD. The data shown is mean ± SD (n ≥ 20) of. The scale bar represents 1 mm. In Figs. b and d, different lowercase letters, shown above the bars, denote statistically significant differences between the samples (Two-way ANOVA followed by Tukey’s post-hoc test, p < 0.05). Numbers indicate fold changes between 22°C and 27°C. **(E** and **F)** Immunoblot analysis of PIF4-FLAG protein levels in the indicated genotypes grown for six days at 22°C and 27°C, and samples were harvested at ZT23 (e), or 22°C SD grown five-day-old seedlings shifted to 27°C for 24h and sampled at ZT23 (f). **(G)** Immunoblot showing PIF4-FLAG protein levels in the indicated genotypes grown under 22°C SD for six days were shifted to 27°C for 24 h, followed by MG132 treatment for 16 h before tissues were harvested. In Figs. **E-G**, Col-0 was used as a negative control. The relative PIF4 protein levels were normalized against Actin and are shown underneath each immunoblot. The normalized PIF4-FLAG protein levels in PIF4-OE at 22°C short day conditions were set to 1, and others were calculated as relative levels. We detected two bands of PIF4-FLAG, and the top band probably corresponds to the phosphorylated form of PIF4. Numbers indicate fold changes between 22°C and 27°C. **(H)** Immunoblots of coimmunoprecipitation assays showing BBX24 interacts with PIF4 in planta. Total protein from six-day-old Col-0 and *35S::BBX24-GFP* seedlings grown at 27°C, sampled at ZT2, were used, and BBX24-GFP was pulled down using anti-GFP antibody coupled to magnetic Dynabeads. BBX24-GFP and PIF4 from input and pulldown fractions were detected by immunoblot using anti-GFP and native anti-PIF4 antibodies. Asterisk denote a non-specific band. **(I)** Immunoblots of coimmunoprecipitation assay showing BBX25 interacts with PIF4 in planta. Total protein from six-day-old Col-0 and *35S::Myc-BBX25* seedlings grown at 27°C, sampled at ZT2, were used, and Myc-BBX25 was immunoprecipitated using an anti-myc antibody coupled to magnetic Dynabeads. Myc-BBX25 and PIF4 from input and pulldown fractions were detected by immunoblotting using anti-Myc and native anti-PIF4 antibodies. **(J)** BiFC assays show the physical interaction of BBX24 and BBX25 with PIF4 in planta in onion epidermal cells. The adaxial epidermis of the onion bulb was co-infiltrated with nYFP, BBX24-nYFP, or BBX25-nYFP, and cYFP or PIF4-cYFP as indicated. Two days after incubation, onion peels were analysed for fluorescence in different combinations of co-infiltered samples. **(K)** BBX24 and BBX25 physically interact with PIF4 in yeast. Overnight grown cultures of Y2H Gold yeast strain containing different combinations of bait (pGBKT7, pGBKT7-BBX24, and pGBKT7-BBX25) and prey (pGADT7 and pGADT7-PIF4) constructs diluted to an OD_600_ of 0.5 with 4x serial dilutions were spotted on double and quadruple dropout medium supplemented with X-alpha-gal. Pictures were taken after 72 h of incubation at 30°C. The experiments were repeated at least three times, and similar results were obtained. All the interactions were repeated three times, and similar results were observed.

Notably, in line with the reduced growth and gene expression phenotype, the PIF4 protein levels was moderately reduced in the single mutants but almost undetectable in the double mutant background both at the end of the night (Figure 4E) and during the day (Supplemental Figure S9B). Similarly, when 22°C SD grown seedlings were exposed to short-term warmth (27°C) for 24h, the PIF4 protein levels was markedly reduced in both *bbx24 PIF4-OE* and *bbx25 PIF4-OE* compared to *PIF4-OE* line (Figure 4F). Interestingly, the PIF4 protein was almost undetectable in the *bbx24bbx25 PIF4-OE* mutant (Figure 4F). Consistent with the unstable nature of the PIF4 protein in the *bbx24bbx25* double mutant, we found that the *PIF4* transcript accumulation was drastically reduced in the *bbx24bbx25* double mutant than Col-0, both at 22°C and 27°C (Supplemental Figure S9C). Also, in line with the reduced *PIF4* mRNA accumulation in the *bbx24bbx25* double mutant observed above, treating *bbx24bbx25* double mutants with MG132 only partially stabilized the PIF4 protein (Figure 4G) further confirms that BBX24/BBX25-mediated PIF4 protein accumulation is manifested both at the transcription and posttranslational level.

In line with the role of BBX24/BBX25 in facilitating PIF4 protein accumulation, overexpression of *BBX24* or *BBX25* in the *PIF4-OE* background resulted in further enhancement in the hypocotyl length while they further suppressed the small cotyledon phenotype of *PIF4-OE* in SD (Supplemental Figure S9D-G) and LD (Supplemental Figure S10A-D) conditions. The *PIF4-OE BBX24-OE* and *PIF4-OE BBX25-OE* double overexpression transgenic lines showed etiolated growth with very long hypocotyl and half-opened or closed cotyledons, especially at 27°C, which is a characteristic phenotype of genotypes with very high PIF4 activity (Supplemental Figure S9D and E). In line with the exaggerated growth phenotypes of the double overexpression transgenic lines, immunoblotting data revealed that when 22°C LD grown seedlings were exposed to 27°C for 4h, PIF4 protein levels was elevated in the *PIF4-OE BBX24-OE* and *PIF4-OE BBX25-OE* transgenic seedlings compared to *PIF4*-*OE* (Supplemental Figure S10E). The *PIF4* transcript levels in these *BBX24-OE PIF4-OE* and *BBX25-OE PIF4-OE* transgenic seedlings was significantly more than the Col-0 but lower than the *PIF4-OE* line, both at 22°C and 27°C under SD (Supplemental Figure S10F), which is likely due to feedback regulation to regulate *PIF4* threshold levels.

### BBX24/BBX25 physically interact with PIF4

Next, to know if BBX24/BBX25-mediated PIF4 protein stabilization at the posttranslational level requires direct physical interactions, we carried out a Co-immunoprecipitation (Co-IP) assay. When BBX24-GFP protein was pulled down using an anti-GFP antibody, we could detect PIF4 protein in the immunoprecipitated complex using a native PIF4 antibody (Figure 4H), while in Col-0 (used as a negative control), we didn’t detect PIF4 protein, suggesting that BBX24 physically interacts with PIF4 in vivo (Figure 4H). Similarly, when we pulled down Myc-BBX25 using an anti-cMyc antibody, we could detect the PIF4 protein in the immunoprecipitated complex (Figure 4I) but not in Col-0, implying that BBX25 physically interacts with PIF4 in vivo (Figure 4I). Consistent with Co-IP data, Bimolecular Fluorescent Complementation (BiFC) assays from onion epidermal cells also revealed BBX24 and BBX25 physically interacts with PIF4 in planta (Figure 4J). To further confirm these interactions, we also performed yeast two-hybrid (Y2H) assays by cloning BBX24 and BBX25 with GAL4 DNA-binding domain (DBD) and PIF4 with Activation domain (AD). Our Y2H assay revealed that BBX24/PIF4 and BBX25/PIF4 co-transformed colonies were able to grow on the minimal media lacking Histidine and Adenine and were also able to activate the production of α-galactosidase, which converts 5-Bromo-4-chloro-3-indolyl α-D-galactopyranoside (X-α-Gal) substrate into and blue colour (Figure 4K). Together, these shreds of evidence confirm that BBX24 and BBX25 physically interact with PIF4 in vivo.

### ELF3 inhibits *BBX24*/*BBX25* gene expression during dawn in a temperature-dependent manner

ELF3, ELF4 and LUX ARRHYTHMO (LUX) form the Evening Complex (EC). ELF3 is the key component of EC as it bridges ELF4 and LUX (Nusinow *et al*., 2011) and is a crucial negative regulator of thermosensory signaling (Box *et al*., 2015; Jung *et al*., 2020; Nieto *et al*., 2015). ELF3 strongly inhibits hypocotyl growth under optimal temperatures by directly inhibiting PIF4 transcription and its transcriptional activity through direct physical association and sequestration (Nieto *et al*., 2015; Nusinow *et al*., 2011). Our above results show that *BBX24*/*BBX25* mRNA expression peaks at the beginning of the day and is drastically reduced during the end of the day through to early night. However, their expression was significantly elevated under warm temperatures, even during the evening, as shown above (Figure 2A). To know if ELF3 inhibits *BBX24/BBX25* gene expression in the evening, we analyzed gene expression through qPCR from Ws (Wild-type) and *elf3* mutant seedlings harvested at ZT6 and ZT10 grown in SD. The expression of *BBX24* and *BBX25* was comparable between Ws and *elf3* mutant at ZT6 (Figure 5A and 5B). Interestingly, their expression is significantly increased in *elf3* mutant at ZT10 compared to Ws, both at 22°C and 27°C (Figure 5A and 5B), suggesting the plausible role of ELF3 in inhibiting *BBX24/BBX25* gene expression during the evening when the EC activity peaks.

**Figure 5.**
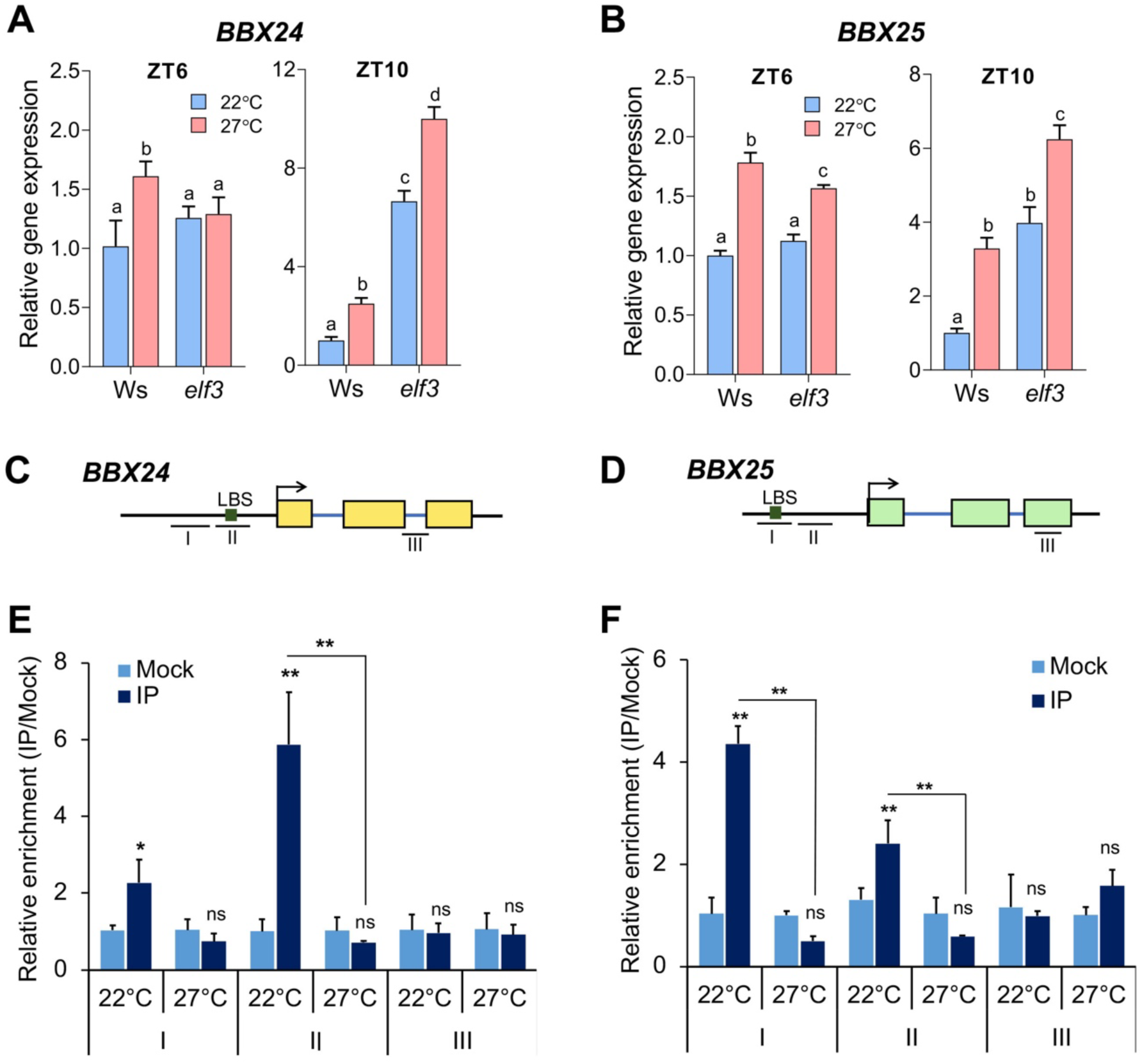
ELF3 represses *BBX24*/*BBX25* transcription during dawn while warm temperature dampens it. **(A** and **B)** qRT-PCR analysis showing relative transcript levels of *BBX24* and *BBX25* in six-day-old seedlings of Ws and *elf3* grown at constant 22°C and 27°C SD conditions, sampled at ZT6 and ZT10. The expression level of each sample was normalized to Ws at 22°C of the respective time point, *and EF1α* was used as an internal control. The data shown is the mean ± SD, with Error bars depicting SD (n=3 biological replicates). **(C** and **D)** Schematic representation of *BBX24* and *BBX25* promoters depicting LUX binding sites and the exact position of each PCR fragment used for analysis. **(E** and **F)** ChIP-qPCR assay showing ELF3 enrichment onto *BBX24* and *BBX25* promoters in vivo. Six-day-old seedlings of *35S::ELF3-YFP* (*elf3-4*) grown at 22°C and 27°C SD conditions were sampled at ZT10 for cross-linking. For each amplicon, fold enrichment was calculated as a ratio between anti-GFP and mock (pre-immune serum). Genomic fragment towards the *3’UTR* of *BBX24* and *BBX25* was used as a negative control. The data shown is the (mean ± SD, n=2, two biological replicates. The experiment was repeated twice, and similar results were seen. Asterisks denote statistically significant differences as determined by Student’s t-test, *P<0.05, **P<0.01.

LUX is a DNA-binding component of EC and has been shown to bind to conserved cis-acting variable motifs such as *A<u>GATTCG</u>A*, *A<u>GATACG</u>C*, *A<u>TATTCG</u>A*, *<u>AAGATC</u>TT* and *G<u>GATCC</u>GA* (Chow et al., 2012; Helfer et al., 2011; Nusinow *et al*., 2011; Silva et al., 2020). We looked for potential LUX binding sites to further understand if ELF3 (a component of EC) can directly bind to *BBX24* and *BBX25* promoters. Interestingly, the *BBX24* and *BBX25* promoters harbour potential LUX binding motifs, *<u>GATACG</u>* and *<u>AAGATC</u>G*, respectively (Figure 5C and D), which are similar to the previously reported motifs bound by LUX (Silva *et al*., 2020). We carried out a Chromatin Immunoprecipitation (ChIP) assay using a *35S::ELF3-YFP* transgenic line expressing stable protein at ZT10 under 22°C and 27°C under SD (Supplemental Figure 11A). Our ChIP qPCR results revealed a significant enrichment of ELF3 onto the *BBX24* and *BBX25* promoters at the region containing the LUX-Binding site in the IP samples compared to mock (preimmune serum) (Figure 5E and 5F). However, the binding of ELF3 to *BBX24* and *BBX25* promoters was strongly inhibited at 27°C grown seedlings (Figure 5E and 5F). This is also consistent with the earlier reports that EC binding to the target DNA was compromised at warm temperatures (27°C) (Silva *et al*., 2020). In summary, our study demonstrates *BBX24*/*BBX25* as novel targets of EC for transcriptional repression at lower temperatures to inhibit growth, while warm temperature-mediated inhibition of ELF3 activity relieves this repression.

### BBX24/BBX25 genetically act downstream to ELF3

Our above results suggest that BBX24/BBX25 have an impact on PIF4 at both protein and transcript levels. ELF3, a potent negative regulator of PIF4, inhibits *PIF4* transcription and also forms heterodimers and sequesters it (Box *et al*., 2015; Nieto *et al*., 2015; Nusinow *et al*., 2011). As ELF3 binds to the *BBX24*/*BBX25* promoter to suppress their gene expression, we hypothesize that ELF3 could be the possible target through which BBX24/BBX25 regulates PIF4 functions. To address this, we performed epistatic interactions by generating *elf3bbx24*, *elf3bbx25* and *elf3bbx24bbx25* mutants. Measurement of hypocotyl length of double, triple and parental genotypes along with WT (segregated from Col-0xWs) suggests that while *bbx24* moderately suppressed, *bbx25* showed weaker suppression of long hypocotyl phenotype (Supplemental Figure S11B and C) and smaller cotyledon area of *elf3* mutant (Supplemental Figure S11D and E). However, *bbx24bbx25* could strongly suppress the long hypocotyl phenotype of *elf3* mutant, as *bbx24bbx25elf3* hypocotyl length was significantly shorter (Figure 6A and 6B). In contrast, the cotyledon area of *bbx24bbx25elf3* was significantly bigger than *elf3* mutant (Supplemental Figure S11F and G), suggesting that *elf3* growth phenotypes are strongly dependent on BBX24/BBX25.

**Figure 6.**
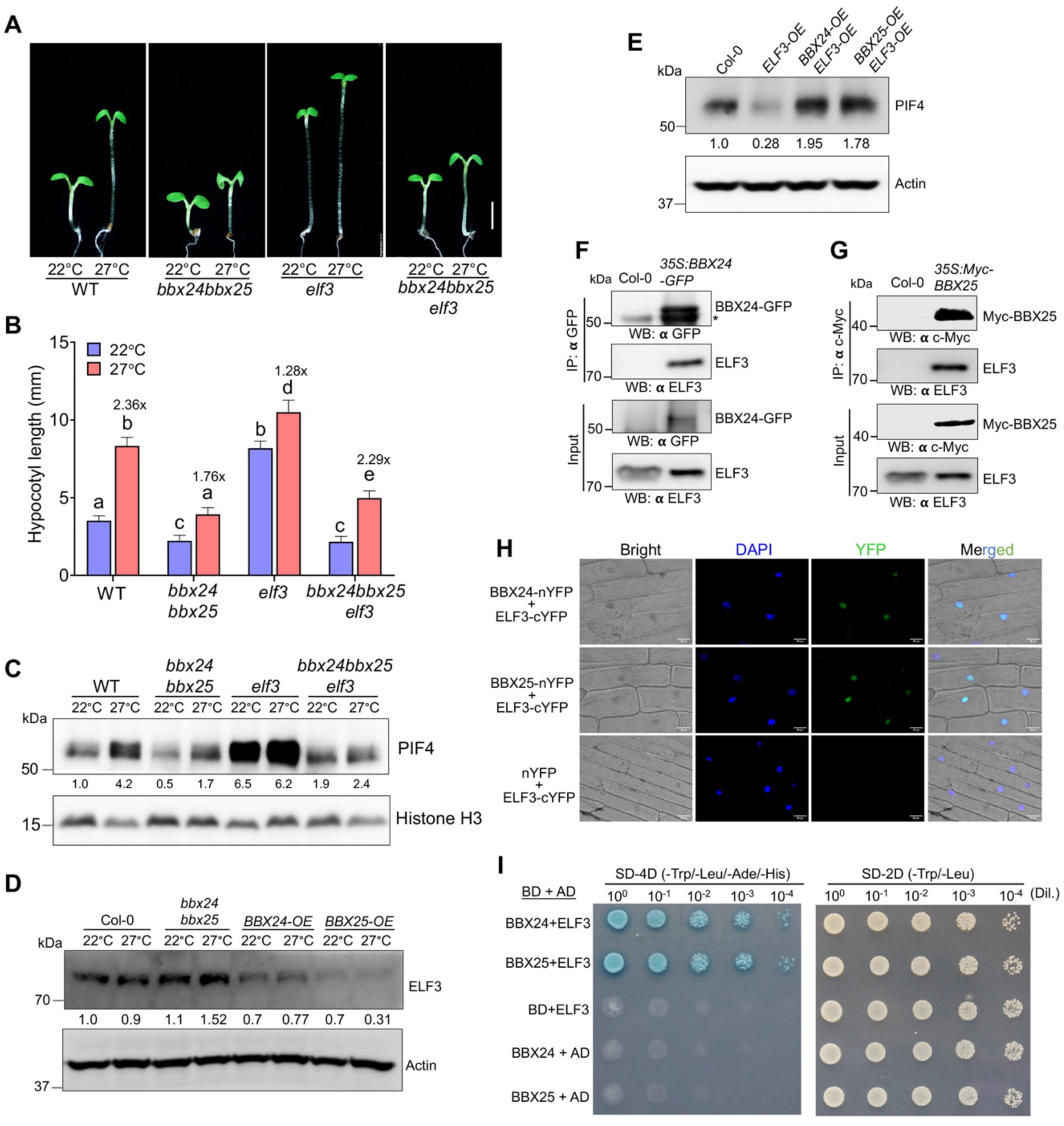
BBX24/BBX25 antagonizes ELF3 to promote PIF4-mediated thermosensory growth. **(A** and **B)** Representative seedling images and hypocotyl length of six-day-old segregated WT (Col-0xWs), *bbx24bbx25*, *elf3* and *bbx24bbx25elf3* seedlings grown at 22°C and 27°C under SD. The scale bar = 2 mm. In Figure b, different lowercase letters, shown above the bars, denote statistically significant differences between the samples (Two-way ANOVA followed by Tukey’s post-hoc test, p < 0.05). Numbers indicate fold changes between 22°C and 27°C. **(C)** Immunoblots showing PIF4 protein levels in WT (Col-0xWs), *bbx24bbx25*, *elf3* and *bbx24bbx25elf3* seedlings grown at 22°C for six days or transferred to 27°C for 24 hours and harvested at ZT23. Histone H3 was used as a loading control, and the relative levels of PIF4 protein normalized to Histone H3 are shown underneath each immunoblot. **(D)** Immunoblot showing ELF3 protein levels in six-day-old Col-0, *bbx24bbx25*, *BBX24-OE* and BBX25-OE seedlings grown at constant 22°C and 27°C under short-day conditions. Actin was used as a loading control, and the relative ELF3 protein level was calculated against Col-0 at 22°C SD. **(E)** Immunoblot analysis of PIF4 protein levels of the indicated genotypes was shifted to 27°C for 24 hours after growing for five days at 22°C SD conditions. Actin was used as a loading control, and the relative PIF4 protein levels shown underneath the immunoblot were calculated against Col-0. **(F** and **G)** Co-immunoprecipitation assay showing in vivo protein-protein interactions between BBX24-ELF3 and BBX25-ELF3. Total protein from six-day-old Col-0, *35S:BBX24-GFP* and *35S:Myc-BBX25* seedlings grown at 22°C SD, harvested at ZT8, were used. The BBX24-GFP and BBX25-Myc proteins were pulled down using anti-GFP and anti-Myc antibodies coupled to magnetic Dynabeads. In the input and immunoprecipitated protein complex, ELF3 was detected using native anti-ELF3 antibodies. Similarly, all the proteins were detected using respective antibodies in the input and immunoprecipitated fractions. Asterisk denote a non-specific band. **(H)** BiFC assays show the physical association of BBX24 and BBX25 with ELF3 in onion epidermal cells. The adaxial epidermis of the onion bulb was co-infiltrated with nYFP, BBX24-nYFP or BBX25-nYFP with cYFP or ELF3-cYFP, as indicated. After two days of incubation, onion peels were observed for fluorescence under a microscope in different co-infiltrate samples. Scale bar = 50 µm. **(I)** Yeast two-hybrid assay showing BBX24 and BBX25 physical interaction with ELF3 in yeast. Y2H Gold strain harbouring different combinations of bait (pGBKT7, pGBKT7-BBX24, and pGBKT7-BBX25) and prey (pGADT7 and pGADT7-ELF3) constructs were grown overnight to an OD_600_ of 0.5 with 4x serial dilutions were dropped on double and quadruple dropout medium supplemented with X-alpha-gal. Pictures were captured after incubating the plates for three days at 30°C. The experiments were repeated at least three times, and similar results were obtained.

As our genetic and molecular data reveal that BBX24/BBX25 probably function downstream to ELF3 but upstream to PIF4, we wanted to see if the suppression of *elf3* growth phenotype by *bbx24bbx25* is due to reduced PIF4 protein levels. Interestingly, our immunoblot data from six-days-old 22°C SD-grown seedlings or 22°C SD-grown seedlings exposed to 27°C for 24h indeed showed that PIF4 protein levels was markedly reduced in the *bbx24bbx25elf3* triple mutant compared to *elf3* mutant both at 22°C and 27°C (Figure 6C). Moreover, increased PIF4 accumulation in *elf3* mutant at ZT23 and ZT8 in 22°C and 27°C grown seedlings was found to be dependent on BBX24/BBX25, as PIF4 protein levels was strikingly reduced in the *bbx24bbx25elf3* triple mutant (Supplemental Figure 12A). The increased PIF4 protein levels in the *elf3* mutant is likely due to increased accumulation of *PIF4* transcript. In line with this, we found that the *PIF4* transcript levels was significantly reduced in the *bbx24bbx25elf3* triple mutant compared to *elf3* (Supplemental Figure S12B). Consistent with the suppression of *elf3* phenotype by *bbx24bbx25*, the expression of PIF4-regulated growth genes such as *XTR7*, *IAA19*, *PRE5,* and *IAA29* are significantly downregulated in the *bbx24bbx25elf3* mutant background than *elf3* (Supplemental Figure S12C). Together, these results confirm that BBX24/BBX25-mediated inhibition of ELF3 function is necessary for optimal PIF4 gene expression and protein accumulation.

Consistent with the role of BBX24/BBX25 in thermosensory hypocotyl and cotyledon growth, we also found that they play an important role in later developmental stages, such as petiole elongation and flowering. We found that long petiole and early flowering phenotypes of *PIF4-OE* and *elf3* mutants were strongly suppressed by *bbx24bbx25* mutations both under SD (Supplemental Figure S13) and LD (Supplemental Figure S14) conditions.

### BBX24/BBX25 antagonizes ELF3 to facilitate PIF4 function

As our above data suggest that BBX24/BBX25 functions antagonistic to ELF3, we hypothesized that BBX24/BBX25 might inhibit ELF3 activity to promote PIF4 functions. To further test this, we introduced *BBX24-OE* and *BBX25-OE* transgenes into the *ELF3-OE* background, generated *BBX24-OE ELF3-OE* and *BBX25-OE ELF3-OE* double homozygous transgenic lines and analyzed their hypocotyl phenotype. Measurement of hypocotyl length from SD-grown seedlings at 22°C and 27°C revealed that overexpression of *BBX24* or *BBX25* significantly suppressed the short hypocotyl phenotype of the *ELF3-OE* transgenic line (Supplemental Figure S15A and B). Consistent with this, our immunoblot data using ELF3 specific antibody (Supplemental Figure S15D) revealed that ELF3 protein level is elevated in the *bbx24bbx25* double mutant but markedly reduced in the *BBX24* and *BBX25* overexpression lines compared to Col-0 (Figure 6D). In contrast to the effect of *bbx24bbx25* mutations in increasing ELF3 protein levels, *ELF3* gene expression was not altered in the *bbx24bbx25* double mutant compared to Col-0 (Supplemental Figure 15C). Further, consistent with the suppression of short hypocotyl length of *ELF3-OE* line by *BBX24-OE* and *BBX25-OE*, we found that, overexpression of *BBX24* and *BBX25* strongly recovered the reduced PIF4 protein levels of *ELF3-OE* line (Figure 6E) indicating negative inhibitory effects of BBX24/BBX25 on ELF3 to facilitate PIF4 protein accumulation.

Next, to know if BBX24/BBX25-mediated inhibition of ELF3 function requires direct physical interactions, we carried out a Co-IP assay. We used *35S:BBX24-GFP* transgenic lines and pulled down BBX24-GFP from the total protein extract using an anti-GFP antibody. In the BBX24-GFP immunoprecipitated complex, we could detect ELF3 as revealed by immunoblotting using a native anti-ELF3 antibody, but not in the Col-0 (Figure 6F). Similarly, when we pulled down from the total protein extract of *35S:Myc-BBX25* transgenic line, we could detect ELF3 in the immunoprecipitated complex but not in Col-0 (Figure 6G), indicating that BBX24 and BBX25 physically interact with ELF3 in vivo. To support these physical interactions, we performed a BiFC assay and found that BBX24 and BBX25 physically interact with ELF3 in planta in onion epidermal cells (Figure 6H, Supplemental Figure S15E). When we co-transformed BBX24-nYFP/ELF3-cYFP and BBX25-nYFP/ELF3-cYFP constructs, we observed reconstitution of uniform YFP fluorescence in the nucleus (Figure 6H). Interestingly, we also observed two or more discrete condensates in the nucleus (Figure 6H; Supplemental Figure 15F). Furthermore, Y2H assays, using ELF3 as bait and BBX24 and BBX25 as prey, found that BBX24 and BBX25 physically interact with ELF3 (Figure 6I). The BD-BBX24/AD-ELF3 and BD-BBX25/AD-ELF3 combinations of constructs, when transformed onto Y2H-Gold reporter strain, they grew on the synthetic minimal media lacking Histidine and Adenine and also, these colonies developed blue colour in the presence of X-α-Gal substrate (Figure 6I). Together, our genetic and biochemical data confirm that BBX24/BBX25 inhibit ELF3 function to promote PIF4-mediated thermomorphogenic growth.

## DISCUSSION

Rising global temperatures due to climate change profoundly affect plant health and pose a significant threat to crop yields (Burroughs *et al*., 2023; Koester *et al*., 2014; Peng *et al*., 2004; Zhao *et al*., 2017). While plants have evolved intricate signaling mechanisms to sense, integrate, and adapt to temperature fluctuations, the underlying signaling mechanisms are poorly understood. As PIF4 is the central component of thermomorphogenesis, maintaining its protein abundance is crucial in achieving optimal thermomorphogenic growth both during the night and daytime. BBX proteins have emerged as crucial signaling nodes for orchestrating adaptive responses to diverse environmental signals such as light, shade, and abiotic stresses (Ding *et al*., 2018; Gangappa and Botto, 2014; Podolec et al., 2022; Song *et al*., 2020; Tripathi et al., 2017; Vaishak *et al*., 2019; Wang et al., 2014; Zhao et al., 2020). This study discovers that BBX24/BBX25 are novel and essential components of the thermosensory signaling pathway and act to promote PIF4 function by inhibiting its critical repressor, ELF3 (Figure 7).

**Figure 7.**
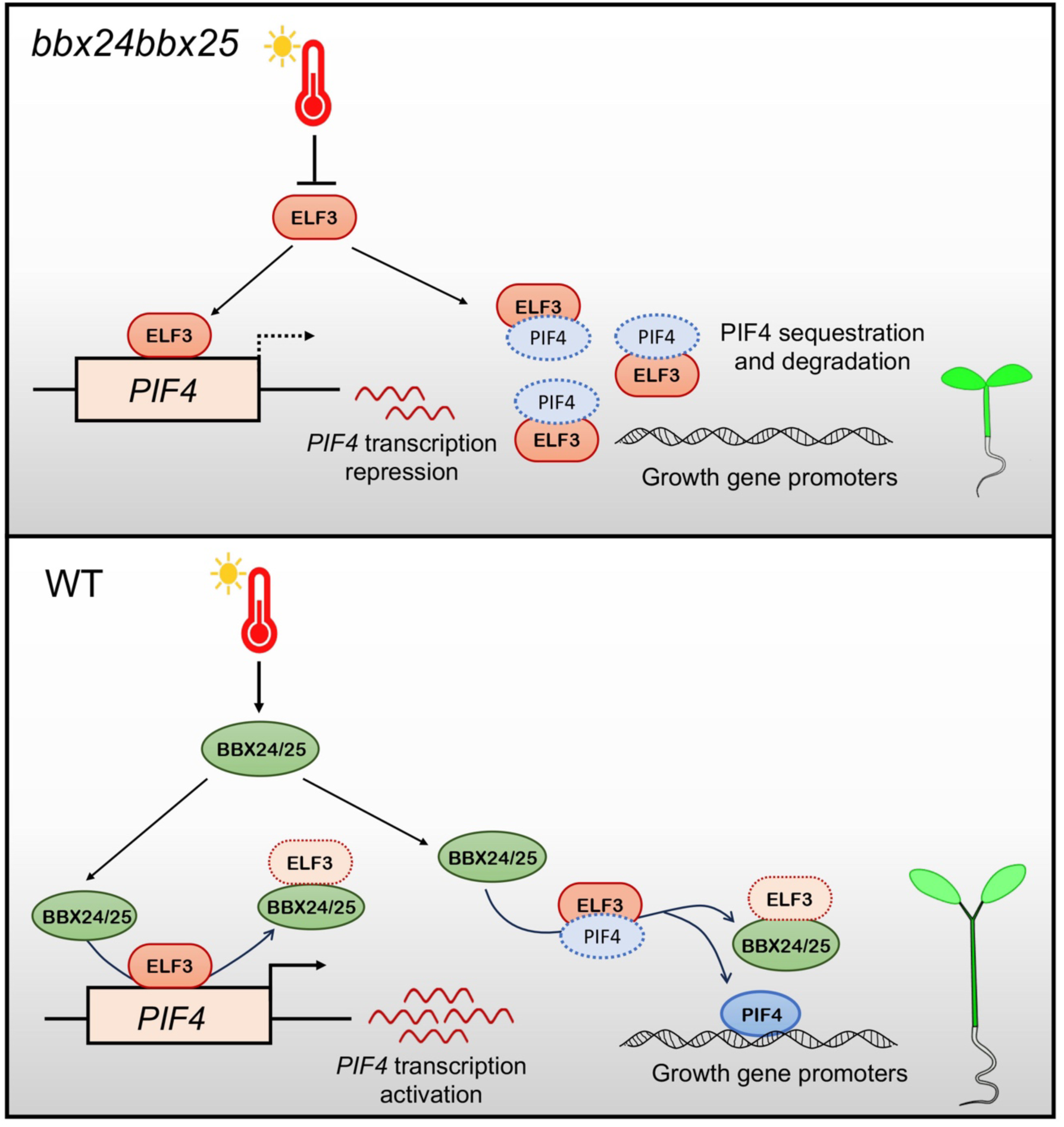
The molecular model depicting BBX24/BBX25-mediated regulation of thermosensory growth. The model elucidates the molecular mechanism by which BBX24/BBX25 promotes thermosensory growth in Arabidopsis. In the absence of BBX24/BBX25, ELF3 exerts a robust inhibitory effect on PIF4 at the transcriptional and protein activity levels, resulting in compromised thermosensory responses. However, in the presence of BBX24/BBX25, they antagonize ELF3, relieving its inhibitory effect on PIF4, resulting in enhanced PIF4 function, and promoting warm temperature-mediated growth.

Our study demonstrates that BBX24/BBX25 play a positive regulatory role in temperature-responsive hypocotyl growth (Figure 1). BBX24/BBX25 functions exhibit significant overlap, as evidenced by a more pronounced reduction in cell length and hypocotyl growth in the *bbx24bbx25* double mutant than in the *bbx24* and *bbx25* single mutants at warm temperature, as opposed to *BBX24-OE* and *BBX25-OE* overexpression lines that promoted cell elongation and hypocotyl growth (Figure 1). Consistent with the growth phenotypes, *bbx24bbx25* double mutants had reduced expression of PIF4 target genes such as *XTR7*, *YUC8*, *EXP8*, *IAA19*, etc., involved in hormone biosynthesis and signaling both at night and during the day (Figure 1; Supplemental Figure S4), which is in line with the reduced PIF4 protein accumulation in response to warm temperatures (Figure 3). Contrary to *bbx24bbx25* mutants, the enhanced thermomorphogenic response and gene expression of the *BBX24-OE* and *BBX25-OE* lines, was consistent with the increased PIF4 protein levels (Figure 3C). Our genetic analysis demonstrates that BBX24/BBX25-mediated thermomorphogenic response is directed through PIF4 (Supplemental Figure S7). In agreement with the BBX24/BBX25 acting through PIF4, the genetic analysis also revealed that the absence of *BBX24* and *BBX25* strongly suppress the robust growth and gene expression phenotypes of the transgenic line overexpressing *PIF4-FLAG*, which is due to the reduced PIF4-FLAG protein accumulation in the single mutants, while complete absence of PIF4-FLAG protein in the *bbx24bbx25* background (Figure 4). The lack of PIF4-FLAG protein was somewhat surprising compared to native PIF4 protein. The possible reason could be the enhanced degradation of PIF4-FLAG protein over native PIF4 protein.

Several thermosensory signaling components such as HMR (HEMERA), RCB (REGULATOR OF CHLOROPLAST BIOGENESIS), and HSFA1s (HEAT SHOCK TRANSCRIPTION FACTOR 1) have been shown to stabilize PIF4 and promote daytime mediated thermomorphogenic growth (Qiu *et al*., 2021; Tan et al., 2023b). Meanwhile, COP1/DET1/SPA, upstream regulators of PIF4, promote PIF4 protein accumulation during the night (Gangappa and Kumar, 2017). Our study reveals that BBX24/BBX25 serve as novel upstream regulators, essential for maintaining optimal PIF4 levels at night and also during the day to promote thermomorphogenesis (Figure 1 and 3). As BBX24/BBX25 and COP1/DET1 promote PIF4 accumulation, their possible functional connection in maintaining optimal PIF4 protein levels in the dark period needs further investigation. While BBX24/BBX25 largely contribute to PIF4 protein accumulation by enhancing *PIF4* gene expression, likely, they are also protecting PIF4 protein from its negative regulators at the post posttranslational level, as revealed from our MG132 and CHX treatment experiments (Figure 3E-3H). The physical interactions between BBX24/BBX25 and PIF4 (Figure 4) also provide evidence that these proteins likely play a crucial role in safeguarding PIF4 from its repressors.

Previous studies have shown that ELF3 acts upstream of PIF4 and inhibits its function both at the transcription and at the protein level (Box *et al*., 2015; Nieto *et al*., 2015; Nusinow *et al*., 2011). Interestingly, our data demonstrate that the *elf3* mutant growth phenotypes, gene expression, and PIF4 protein accumulation strongly depend on BBX24/BBX25, suggesting that BBX24/BBX25 likely act downstream to ELF3 but upstream to PIF4 (Figure 6). Our RT-qPCR and ChIP data further support that ELF3 inhibits *BBX24* and *BBX25* gene expression during the evening at 22°C, which is nullified at 27°C (Figure 5), due to warm temperature-mediated inactivation of ELF3 activity (Jung *et al*., 2020). Nevertheless, our immunoblot data revealed that BBX24/BBX25 inhibit ELF3 protein accumulation at warm temperatures (Figure 6D). Therefore, in addition to warm temperature-mediated inhibition of ELF3 activity through liquid-liquid phase separation (LLPS), BBX24/BBX25 inhibit ELF3 function by reducing ELF3 accumulation. Moreover, the physical association of BBX24/BBX25 could also sequester with ELF3, resulting in enhanced PIF4 function (Figure 6). This is also supported by genetic data, wherein *BBX24-OE* and *BBX25-OE* significantly suppressed the short hypocotyl phenotype and reduced PIF4 accumulation in the *ELF3-OE* transgenic line (Figure 6E; Supplemental Figure S15A and B). Therefore, we propose that constitutive thermomorphogenic response and high PIF4 activity in the *elf3* mutant is not solely attributed to the alleviation of transcriptional repression on PIF4 but also due to the concurrent increase in BBX24/BBX25 activity, and these dual mechanisms together potentiate thermomorphogenic response (Figure 6; Supplemental Figure S11 and S12).

Consistent with the role of BBX24/BBX25 in promoting PIF4 function, the PIF4-mediated thermosensory petiole growth and flowering are also dependent on PIF4 both under SD and LD photoperiods, highlighting their role beyond seedling development (Supplemental Figure S13 and S14). These pieces of evidence reveal that BBX24/BBX25 proteins act as key regulators of growth and development in response to varying light and temperature cues by coordinating clock and external cues. In summary, we propose the ELF3-BBX24/BBX25-PIF4 as a novel regulatory module (Figure 7), critical for integrating temperature and light cues to regulate thermosensory growth and development in Arabidopsis.

## METHODS

### Plant Material and Growth Conditions

All the lines used here are in Columbia-0 (Col-0) background except *elf3*-4, which is in Wassilewskija (WS) background. The genotypes *bbx24-1* (Gangappa *et al*., 2013), *bbx25-2* (Gangappa *et al*., 2013) *35S::BBX24-GFP*(Job *et al*., 2018), *pPIF4*::*PIF4-FLAG* (Gangappa *et al*., 2017), *pif4-101* (Lorrain *et al*., 2008), and elf3-4 {Dixon, 2011 #2021), have been described previously. For hypocotyl experiments, seeds were surface sterilized with 70% ethanol containing 0.5% Triton X-100 for 10 minutes and plated on half-strength Murashige and Skoog (MS) medium (Sigma-Aldrich) supplemented with 1% sucrose, 0.5 mM MES (pH-5.7) and 0.8% agar following stratification at 4°C for four-days. For the hormone experiments, seeds were plated on MS media containing different concentrations of hormones or their inhibitors. Upon seed germination at 22°C long day (16h Light/8h Dark) growth chamber (Percival Scientific) seedlings were transferred to respective 22°C and 27°C short day (8h Light/16h Dark) conditions or as stated in the experimental conditions with 70% humidity and light intensity of 100 µmol m^−2^ sec^−1^. Five days after germination (six-day-old), seedlings were taken for the hypocotyl length and cotyledon area measurements using ImageJ software. At least 25 seedlings for each genotype were taken for the measurements.

### Generation of *BBX25* overexpression transgenic lines

For the generation of *35S::Myc-BBX25* overexpressing transgenic lines, the full-length BBX25 coding sequence was amplified from 22°C LD grown Col-0 cDNA and cloned into the pENTR vector through pENTR™ Directional TOPO^®^ Cloning Kits (Invitrogen), which is then subcloned into the destination vector pGWB618 (Nakamura et al., 2010) using LR cloning kit (Invitrogen). The binary expression cassette is then transformed into *Agrobacterium tumefaciens* GV3101 strain. Agrobacterium harbouring the transgene were transfected into Col-0 through the floral dip method. Positive transformants were selected on ½ MS containing 7.5 ug/ml glufosinate-ammonium (BASTA). Four independent lines showing 3:1 segregation on BASTA were chosen for further analysis. In the T_3_ generation, these lines were selected based on transgene and protein expression analysis for further analysis.

### Generation of double and triple mutants

For the generation of double mutants *bbx24-1bbx25-2*, *bbx24-1 PIF4-OE*, *bbx25-2 PIF4-OE*, *BBX24OE pif4-101*, *BBX25-OE pif4-101*, *bbx24-1 elf3-4*, *bbx25-2 elf3-4*, *BBX24-OE PIF4-OE*, *BBX25-OE PIF4-OE*, *BBX24-OE ELF3-OE* and *BBX25-OE ELF3-OE* we crossed *bbx24-1* and *bbx25-2* with each other and *bbx24-1*, *bbx25-2*, *BBX24-OE* and *BBX25-OE* with the respective genotypes. For the generation of triple mutants *bbx24-1 bbx25-2 PIF4-OE* and *bbx24-1 bbx25-2 elf3-4* we crossed homozygous double mutant of *bbx24-1 PIF4-FLAG* with *bbx25-2 PIF4-FLAG* and *bbx24-1elf3-4* with *bbx25-2elf3-4*, respectively. After crossing, we got F_1_ seeds that grew for one more generation to get F_2_ seeds. Approximately 20-30 F_2_ seedlings growing on antibiotic selection were picked up where homozygosity was confirmed through PCR-based genotyping method using gene-specific and T-DNA specific primers. T-DNA-specific primers used were SALK LB1.3 for *bbx24-1*, SAIL LB3 for *pif4-101*, and *spm1* for *bbx25-2*. For *elf3-4* genotyping, we amplified PCR fragments using dCAPS primers followed by AluI restriction digestion. To confirm the homozygosity of *BBX24-OE*, *BBX25-OE*, *PIF4-FLAG* and *ELF3-OE*, we first confirmed the presence of CDS using PCR, followed by the survival of seedlings on an antibiotic selection plate in the next generation. F_3_ generation seedlings showing 100% survival on double antibiotic selection were used for further experiments. Primers used for genotyping are listed in Table S1.

### Gene expression analysis

Approximately 100 mg seedlings from the indicated genotypes and growth conditions were harvested, frozen in liquid nitrogen, and stored at −80°C until further processing. For RNA extraction, seedlings were ground to a fine powder in liquid nitrogen, and total RNA was isolated using an RNeasy plant mini kit (Qiagen) following the kit protocol with on-column DNase I digestion (Qiagen) for 30 min. at RT. 1.5 µg of total RNA was converted into cDNA using SuperScript® III First-Strand Synthesis kit (Invitrogen) by taking Oligo(dT)_20_ primers following manufacturer’s instructions. For RT-qPCR, nuclease-free water-diluted cDNA was mixed with PowerUp™ SYBR™ Green Master Mix (ThermoFisher Scientific) and gene-specific primers. Each gene RT-qPCR reaction was set in triplicate with a Biorad CFX96 Touch Real-Time PCR Detection System or an Applied Biosystems^TM^ 7500 Fast Real-Time PCR System. The transcript level of each gene was calculated relative to that of EF1α. RT-qPCR results were analyzed in fold change using the 2^−ΔΔCt^ method.

### Protein extraction and Immunoblotting

For western blotting, approximately 100-150 mg tissue of six-day-old *Arabidopsis* seedlings grown in different experimental conditions were harvested in a microcentrifuge tube, snap-frozen and ground to fine powder in liquid nitrogen. Total protein was extracted in 250 µl of protein extraction buffer containing 50 mM Tris-HCl pH 8.0, 150 mM NaCl, 10% glycerol, 5mM DTT, 1% NP-40, 0.5mM PMSF, and 1X EDTA free Protease Inhibitor Cocktail (Roche). After resuspending in the extraction buffer, crude protein samples were cleared by centrifuging at 15,000 rpm for 10 min at 4°C. Protein concentration was estimated using Bradford reagent (Biorad). Approximately 30 µg of total protein was boiled at 95°C for 5 minutes in 5x SDS gel loading dye (100 mM Tris-HCL pH 7.5, 12% glycerol, 4% SDS, 200 mM DTT, 40 mM β-ME and 0.001% bromophenol blue), separated via SDS-PAGE and transferred to the PVDF membrane (Millipore, Sigma) by wet transfer method.

For protein detection, blots were blocked with 5% skim milk in TBST for 2h at RT, incubated overnight at 4°C with respective primary antibodies, followed by incubation in 1:10,000 dilutions of horseradish peroxidase-conjugated rabbit anti-goat (Thermo Fisher Scientific, 811620) or goat anti-rabbit (Abcam, ab205718) secondary antibodies at room temperature for 1 h. Primary antibodies used were polyclonal goat anti-PIF4 antibodies (Agrisera, AS16 3955, 1:2500), polyclonal rabbit anti-GFP antibodies 1:7500 (Abcam, ab290), monoclonal rabbit anti-GFP (Invitrogen G10362, 1:500) monoclonal HRP conjugated mouse anti-myc (HRP) antibodies (Abcam, ab62928, 1:5000), monoclonal HRP conjugated mouse anti-FLAG antibodies (Abcam, ab49763, 1:5000), polyclonal rabbit anti-HY5 antibodies (Agrisera, AS12 1867, 1:2500), polyclonal rabbit anti-ELF3 antibodies (Agrisera, AS18 4168, 1:2000) polyclonal rabbit anti-actin antibodies (Abcam, ab197345, 1:10,000), and Rabbit polyclonal anti-Histone H3 antibodies, 1:10,000 (abcam, ab1791). The chemiluminescence signal was detected using the SuperSignal West Femto kit (Thermo Fisher Scientific) and Syngene gel-doc system. Protein band intensities were measured using the ImageJ software (https://imagej.nih.gov/).

### MG132 and CHX treatment

Mg132 and CHX treatments were carried out as reported (Zhou et al., 2021) with minor modifications. For MG132 treatment, five-day-old *Arabidopsis* seedlings were transferred to liquid MS medium containing 50µM MG132 or DMSO and then incubated for 8 h or 16 h before tissues were harvested for immunoblot. To determine the degradation rate of PIF4 protein by blocking de novo protein synthesis, five-day-old 22°C SD grown seedlings treated with MG132 as above before shifting to 27°C for 8hrs were washed five times and then transferred to liquid MS medium supplemented with 200µM CHX followed by tissue harvest at different time points for western blot analysis.

### Chromatin Immunoprecipitation (ChIP) assay

ChIP assays were performed as previously described (Yamaguchi et al., 2014) with modifications. Approximately 1.5 g of *35S::ELF3-YFP*(*elf3-4*) seedlings grown for five days after germination at 22°C SD and 27°C SD were harvested and cross-linked with 30 ml of 1% formaldehyde in 1X PBS for 2 × 10 mins by vacuum infiltration. Crosslinking was quenched by adding 125 mM glycine under vacuum for another 5 min. Fixed seedlings were rinsed with 30 ml 1X PBS twice, dried on stacks of filter paper, frozen in liquid nitrogen and stored at −80°C. Frozen samples were crushed in liquid N_2_ until they became light green and homogenized in 30 ml of extraction buffer I (10 Tris-HCL pH 8, 0.4M sucrose, 10 mM MgCl2, 5mM β-Mercaptoethanol, 2 mM PMSF and 1X protease inhibitor cocktail). Lysate was passed through 2 layers of Miracloth to remove debris and centrifuged at 1000 x g for 20 minutes at 4°C to pellet down cell organelles. Pellet resuspended in 5ml of extraction buffer II (10 mM Tris-HCL pH 8, 0.25M sucrose, 10 mM MgCl2, 1% Triton X-100, 5mM β-Mercaptoethanol, 2 mM PMSF and 1X protease inhibitor cocktail) to burst other cell organelles and centrifuged at 1000 x g for 10mins at 4°C. The deposited pellet was then dissolved in 300 µl of extraction buffer III (10 mM Tris-HCL pH 8, 1.7 M sucrose, 2mM MgCl2, 0.15% Triton X-100, 5mM β-Mercaptoethanol, 2mM PMSF and 1X protease inhibitor cocktail) which was carefully layered at the top of 600 μl extraction buffer III and centrifuged at 16,000 x g for 1hr at 4°C to concentrate nuclei. The nuclear pellet was dissolved in 300 μl of sonication buffer, and chromatin was sheared by sonicating for 20 cycles for 30 sec. ON/OFF. Sonicated samples were cleared by centrifuging at 12000 x g for 10 mins. 1/10^th^ of the sonicated sample was kept at −20°C as input, and the rest was diluted 10-fold in ChIP dilution buffer for immunoprecipitation. Protein-DNA complexes were immunoprecipitated by incubating the diluted chromatin overnight at 4°C with 15 μl of Dynabeads Protein-G (Invitrogen) equilibrated with 2 μg of anti-GFP antibody (Takara Bio, 632381). Beads were washed for 2 × 10 mins in low salt wash buffer, 1 × 10 mins in high salt wash buffer, 1 × 10 min in LiCl wash buffer, and 2 × 10 min in TE buffer. DNA from input and IP were eluted by incubation with 10% chele× 100-resin (Biorad) at 95°C for 10mins followed by 20 μg of Proteinase K treatment at 37°C for 30 min. The supernatant was collected after centrifuging at 16000 x g for 2 min., followed by the precipitation of DNA through the phenol-chloroform extraction method. Precipitated DNA was dissolved in 50 μl of nuclease-free water, and 2 μl was used per reaction for chip-qPCR.

For the ChIP assay of BBX24/BBX25 binding on *PIF4* and its target genes promoter, we have used six-day-old *35S:BBX24-GFP* and *35S:Myc-BBX25* seedlings grown at 27°C and tissues were harvested at ZT2 and ZT4, respectively. Further, we have processed the tissues in a similar way as described above for ELF3 ChIP analysis. Primers used for ChIP-qPCR analysis are listed in Supplemental Table 1.

### Yeast two-hybrid (Y2H) assay

Y2H assay was performed following the Matchmaker® Gold Yeast Two-Hybrid System User Manual (Takara Bio). Briefly, full-length coding sequences of *BBX24* and *BBX25* were cloned into the bait vector pGBKT7 containing DNA-Binding Domain (DBD), while PIF4 and ELF3 were cloned into the prey vector pGADT7 having Activation Domain (AD). For checking the protein-protein interactions, different combinations of bait and prey constructs were co-transformed into Y2H Gold yeast strain (Clontech) using Yeastmaker™ Yeast Transformation kit (Clontech), and transformants were selected on double dropout medium SD/-Leu/-Trp at 30°C for three-days. For confirmation of protein-protein interactions, overnight grown double dropout screened yeast cells were diluted to an OD_600_ of 0.5 with 4X serial dilutions were spotted on quadruple dropout medium SD/-Leu/-Trp/-Ade/-His supplemented with X-alpha-gal. Plates were grown at 30°C for 72 hr before taking the picture.

### Co-Immunoprecipitation

Coimmunoprecipitation assays were performed using Dynabeads™ Protein G Immunoprecipitation Kit (Invitrogen). Approximately 500 mg of six-day-old seedlings grown at 27°C for BBX24/BBX25-PIF4 and at 22°C for BBX24/BBX25-ELF3 immunoprecipitation, harvested at ZT2 and ZT8 were used, respectively. Harvested seedlings were grounded in liquid nitrogen, and the powder was homogenized in 1 ml of co-immunoprecipitation buffer containing 50 mM Tris-HCL pH-8, 100 mM NaCl, 10% glycerol, 2mM DTT, 1mM EDTA, 0.1% NP-40, 1mM PMSF, and 1X protease inhibitor cocktail. Crude protein extract was cleared by two rounds of centrifugation at 13000 rpm for 10 mins at 4°C. An aliquot of 100 μl was boiled in 5X SDS loading dye as an input. The remaining 900 μl protein lysate was incubated with 50 μl Dynabeads (Invitrogen) equilibrated with anti-GFP or anti-cMyc antibody at 4°C for 6 h. Then, beads were washed, and immunoprecipitated proteins were eluted by boiling the beads at 70°C for 10 minutes in 1X Laemmli buffer. Then, input and immunoprecipitated protein samples were subjected to SDS-PAGE for immunodetection. For immunoprecipitation assay of BBX24-GFP/Myc-BBX25-PIF4 polyclonal rabbit anti-GFP (Abcam, ab290, 1:5000), monoclonal mouse anti-myc (Abcam, ab62928, 1:5000) and polyclonal goat anti-PIF4 antibodies (Agrisera AS163955, 1:2500) were used. For BBX24-GFP/Myc-BBX25-ELF3 immunoprecipitation, monoclonal rabbit anti-GFP (Invitrogen G10362, 1:500), monoclonal mouse anti-myc (Abcam, ab62928, 1:5000) and polyclonal rabbit anti-ELF3 antibodies (Agrisera, AS18 4168, 1:2000) were used.

### Differential Interference Contrast (DIC) Microscopy for cell length measurement

For histological analysis, 6-day-old whole seedlings were fixed and cleared using serial dilution of ethanol (15%, 50%, 70%, 96% and 100%), each for 15 min in a 24-well plate. The seedlings were then kept in a cold room at 4° C overnight. On a subsequent day, seedlings were rehydrated in an ethanol gradient (100%, 96%, 70%, 50% and 15%), each step for 15 min., after which those were washed with 0.2 M sodium phosphate buffer (pH=7). Aniline blue (Sigma-Aldrich, Germany) staining solution was prepared as 0.1% w/v stock solution in 0.2 M sodium phosphate buffer.

Seedlings were stained with a 1:20 dilution of aniline blue stock solution with 0.2M sodium phosphate buffer. Samples were incubated overnight at room temperature. Seedlings were mounted onto chloral hydrate solution (chloral hydrate: glycerol: Milli-Q water:: 8:1:2), and hypocotyls were observed using an Olympus IX81 inverted microscope under differential interference contrast optics. Images were acquired with cellSens Standard Software (Olympus).

Micrographs of six seedlings were obtained for each genotype, and their cell length was considered under two different temperature conditions, 22°C and 27°C short-day conditions. The cell length of hypocotyls was defined as the average number of cortical cells crossing the central region of hypocotyls-the number of cells varies with the genotype and the growth condition. Only the cells extending in the non-dividing files were counted.

### Bimolecular Fluorescent Complementation (BiFC) assay

For the BiFC assays, the *CDS* of *BBX24*, *BBX25*, *PIF4* and *ELF3* in the pENTR-D/TOPO vectors have been recombined into pSPYNE-nYFP and pSPYCE-cYFP vectors using LR-Clonase (Invitrogen) to obtain the fusion proteins. *BBX24* and *BBX25* were cloned into the pSPYNE vector, whereas *PIF4* and *ELF3* were cloned in the pSPYCE vector. Sequence-verified constructs were then mobilized into *Agrobacterium tumefaciens* (strain GV3101) with the help of the freeze-thaw method. *Agrobacterium-*mediated *in planta* transient transformation onto onion bulb as reported(Xu et al., 2014) The p19 protein of the tomato bushy stunt virus has been used to subdue gene silencing. For infiltration, nYFP-BBX24/c-YFP-PIF4, nYFP-BBX25/cYFP-PIF4, nYFP-BBX24/c-YFP-ELF3 and nYFP-BBX25/cYFP-ELF3 along with the p19 combination, were used with a final concentration of OD_600_ 0.6: 0.6: 1.0. After infiltration, onion bulbs were kept at 28±1 °C in the dark for 72 h. The infected adaxial epidermis of the bulb scale was peeled and visualized with an SP8 confocal microscope (Leica, Germany) using 40x objective (ex: 488 nm for YFP, ex: 405 nm for DAPI) and deconvoluted using Leica Lightning software. Nuclei were counterstained with DAPI (1 µg/ml). A combination of nYFP-BBX24/cYFP, nYFP-BBX25/cYFP, nYFP/cYFP, nYFP/cYFP-PIF4 and nYFP/cYFP-ELF3 were used as the negative controls. For the BBX24/BBX25-ELF3 BiFC assay, an Olympus IX81 epifluorescence microscope was used, and images were acquired using ImageJ Fiji software.

### Quantification of GFP fluorescence using epifluorescence microscope

GFP fluorescence of BBX24-GFP was captured using an Olympus IX81 inverted microscope equipped with an LED lamp. Approximately ten seedlings were taken for imaging after five days of growth in the respective conditions. The BBX24-GFP fusion protein was excited with a 488 nm light, and the emitted fluorescence signal was detected between 500nm and 561nm range. Images were acquired, and fluorescence intensity was quantified using ImageJ’s Fiji software. The mean fluorescence intensity was calculated by marking the area with a freehand selection tool. Subsequently, relative fluorescence intensity was calculated against 27°C seedlings.

### GUS Histochemical staining

For GUS-staining six-day-old 22°C and 27°C SD grown F1 seedlings of *DR5:GUS, BBX24-OEDR5:GUS* and *BBX25-OEDR5:GUS* seedlings were harvested at ZT2 and fixed in 50mM Sodium phosphate buffer containing 2% formaldehyde and 0.05% Triton X-100. After fixation, the seedlings were appropriately cleaned by washing them three times with 50 mM phosphate buffer. To detect the GUS activity, seedlings were submerged and vacuum infiltrated for 4mins in 0.5 mg/ml 5-bromo-4chloro-3-indoyl glucuronide (X-Gluc) solution containing 50mM sodium phosphate buffer, 5mm Na2-EDTA, 1mM potassium ferrocyanide and 1mM potassium ferricyanide following incubation at 37°C for overnight. The next day, seedlings were destained with 75% and 95% ethanol for 3 hours each at 37°C and observed under a stereomicroscope.

### Quantification and statistical analysis

Hypocotyl length, cotyledon area, and protein band intensities were quantified using ImageJ software (https://imagej.nih.gov/). The statistically significant difference in the data sets was analyzed using GraphPad Prism version 8.0 (https://www.graphpad.com/). One-way ANOVA or Two-way ANOVA Tukey test was used for multiple groups of observations. Students’ t-test was used to determine the significance of two groups of observations. Details of the statistical analysis can be found in the figure legends.

## ACKNOWLEDGEMENTS

We thank Dr. Vinod Kumar, Prof. Salome Prat and Prof. Javier F. Botto, respectively, for the *pPIF4:PIF4-FLAG*, *35S::ELF3-YFP*(*elf3-4*) and *35S::BBX24-GFP* transgenic lines seeds. We thank all the members of Gangappa lab for fruitful discussions. B.C.M., S.S. and V.G. acknowledge the Council of Scientific and Industrial Research, Department of Biotechnology, and University Grants Commission, respectively, for their doctoral fellowships. G.U acknowledges the Postdoctoral Fellowship from IISER Kolkata (PDF/DBS/2021/025) and Science & Engineering Research Board (SERB-PDF/2022/000529). R.C. acknowledges the KVPY Fellowship from the Department of Science and Technology, Govt. of India.

## AUTHOR CONTRIBUTIONS

S.N.G. conceptualized, designed and supervised the study, analyzed data, and wrote the first and subsequent drafts of the manuscript with inputs from all the authors. B.C.M. designed the study, planned and conducted most experiments, collected results, analyzed the data, and helped write the manuscript draft. S.S. generated *elf3* and *DR5:GUS*-related genetic lines, conducted phenotypic analysis, and measured cell length. V.G. assisted in raising *pif4-101*-related genetic lines with phenotypic analysis. R.C. contributed to generating novel genetic materials and conducted experiments. G.U. carried out BiFC experiments. All the authors have read and approved the final version of the manuscript.

## DATA AVAILABILITY

The data presented in this study is available in the main figures and the supplementary material.

## SUPPLEMENTAL INFORMATION

**Supplemental Figure S1.** Molecular and phenotypic analysis of BBX25 overexpression transgenic lines.

**Supplemental Figure S2.** BBX24/BBX25 mediated thermosensory hypocotyl growth is linked to growth-promoting hormones.

**Supplemental Figure S3.** BBX24/BBX25 overexpression lines show differential sensitivity to Auxin and BR inhibitors.

**Supplemental Figure S4.** BBX24/BBX25 regulates the expression of hormone and growth-related genes during the daytime.

**Supplemental Figure S5.** Warm temperatures promote BBX24/BBX25 transcript and protein accumulation.

**Supplemental Figure S6.** ChIP analysis for BBX24/BBX25 enrichment on *PIF4* and its target genes promoter.

**Supplemental Figure S7.** BBX24/BBX25 mediated thermosensory hypocotyl growth depends on PIF4.

**Supplemental Figure S8.** BBX24/BBX25 are essential for PIF4-mediated regulation of hypocotyl and cotyledon growth in response to warm temperatures.

**Supplemental Figure S9.** BBX24/BBX25 functions are required for PIF4-governed hypocotyl and cotyledon growth at higher temperatures.

**Supplemental Figure S10.** Regulation of hypocotyl and cotyledon growth by PIF4 requires BBX24/BBX25 under LD photoperiod.

**Supplemental Figure S11.** ELF3 genetically interacts with BBX24 and BBX25 to control thermosensory hypocotyl and cotyledon growth.

**Supplemental Fi. S12.** BBX24/BBX25 inhibit ELF3 function to promote PIF4-governed thermosensory growth.

**Supplemental Figure S13.** ELF3-BBX24/BBX25-PIF4 module controls the thermosensory flowering and petiole growth under SD photoperiod.

**Supplemental Figure S14.** BBX24/BBX25 is required to manifest petiole growth and flowering of the *PIF4-OE* transgenic line and *elf3* mutants under LD photoperiod.

**Supplemental Fig S15.** BBX24/BBX25 antagonize ELF3 function.

**Supplemental Table S1.** List of oligonucleotides used in this study.

